# Continent-wide drivers of spatial synchrony in age structure across wild great tit populations

**DOI:** 10.1101/2024.05.30.596055

**Authors:** Joe P. Woodman, Stefan J. G. Vriend, Frank Adriaensen, Elena Álvarez, Alexander Artemyev, Emilio Barba, Malcolm D. Burgess, Samuel P. Caro, Laure Cauchard, Anne Charmantier, Ella F. Cole, Niels Dingemanse, Blandine Doligez, Tapio Eeva, Simon R. Evans, Arnaud Grégoire, Marcel Lambrechts, Agu Leivits, András Liker, Erik Matthysen, Markku Orell, John S. Park, Seppo Rytkõnen, Juan Carlos Senar, Gábor Seress, Marta Szulkin, Kees van Oers, Emma Vatka, Marcel E. Visser, Josh A. Firth, Ben C. Sheldon

## Abstract

Spatio-temporal variation in age structure influences population dynamics, yet we have limited understanding of the spatial scale at which its fluctuations are synchronised between populations. Using 32 great tit populations, spanning 3200km and *>*130,000 birds across 67 years, we quantify spatial synchrony in breeding age structure and its drivers. We show that larger clutch sizes, colder winters and summers, and larger beech crops lead to younger populations. We report distant-dependent spatial synchrony of age structure, which is maintained at approximately 650km. Despite covariation with age structure, reproductive and environmental variables do not influence the scale of synchrony, except for a moderate effect of beech masting. We suggest that local ecological and density-dependent dynamics impact how environmental variation interacts with age structure, influencing estimates of the environment’s effect on spatial synchrony. Our analyses demonstrate the operation of synchrony in age structure over large scales, with implications for age-dependent demography in populations.

## 1 Introduction

Age-specificity in individual-level traits means that variation in population age structure can feed through to affect various population processes. For example, variation in age structure influences population-level social functioning (Siracusa et al. 2023; Woodman et al. in press), while population growth rate is affected by variation in age structure (Caswell 2000; Sibly & Hone 2002). Further, the influence of age structure on population vital rates is self-reinforcing, in that when demographic rates change, this affects the number of individuals that are recruited and die. Thus, variation in age structure is fundamentally driven by non-stationarity, which arises intrinsically as population demographic rates vary, but also from environmental variability which affects the number of individuals within different cohorts (Koons et al. 2016; Rollinson et al. 2021). While a considerable amount of research has identified within-population temporal variation in age structure (Coulson et al. 2004, 2005; Gamelon et al. 2016), relatively little is known about the spatial scale at which age structure varies, whether temporal dynamics differ between populations, and what between-population differences in fluctuations in age structure suggest about the drivers of its variation.

Spatial synchrony is the concurrent change in time-varying characteristics of spatially-distinct populations (Bjørnstad et al. 1999a; Liebhold et al. 2004), which operates across many animal populations (Elton 1924; Moran 1953; Wan et al. 2022). Spatial synchrony can be important for population stability (Paradis 1997; Ruxton 1994), but highly synchronous dynamics may impose a risk of species extinction if density crashes occur simultaneously (Heino et al. 1997). Research has identified spatial synchrony in survival (Olmos et al. 2020), body mass (Herfindal et al. 2020), breeding success (Olin et al. 2020; Vriend et al. 2023), phenological timing (Vriend et al. 2023), and population size (Bjørnstad et al. 1999a; Hansen et al. 2020; Koenig 1999), particularly in birds (Mortelliti et al. 2015; Paradis et al. 1999, 2000; Sæther et al. 2007). However, despite the interrelated dynamics between variation in population growth and age structure, there has been little research into spatial synchrony of age structure and the mechanisms that might drive this.

Spatial synchrony arises from three primary mechanisms: dispersal between populations (Kendall et al. 2000; Ripa 2000); interspecific trophic interactions with other organisms that are themselves spatially synchronised (Ims & Andreassen 2000; Jones et al. 2003; Selås 1997); or a common influence on populations from environmental variables that are correlated in space – known as the “Moran effect” (Hudson & Cattadori 1999; Moran 1953; Ranta et al. 1997). Understanding the spatial scale at which population age structure co-fluctuates will provide insight into whether any of these mechanisms drive spatial synchrony in age structure. This is particularly important considering climate change is increasingly impacting wild populations through effects on survival (Parmesan 2006) and other population dynamics (Hansen et al. 2020; Ozgul et al. 2010). Thus, gaining insight of spatial synchrony in age structure due to environmental regulation is relevant for understanding the effects of climate change on natural population dynamics, particularly considering that highly synchronous age structure dynamics might induce simultaneous density crashes and prevent the possibility of demographic rescue (Engen et al. 2002; Mills 2012).

In this study, we assess spatial synchrony of variation in age structure across 32 European great tit populations. We first assess whether fluctuations in age structure are explained by reproductive and environmental factors that vary at different spatial scales. Second, we quantify whether temporal fluctuations in age structure depend on distance between populations, and whether such spatial synchrony is explained by variation in reproductive and environmental variables. By assessing the influence of separate explanatory variables, we identify how aspects of reproductive and environmental variability differentially influence variation in age structure, and their role in synchronising population dynamics.

## 2 Methods

### 2.1 Study system and data collection

The great tit *Parus major* is a passerine bird found in mixed woodlands across much of the Western Palearctic. Their reproductive lifespan ranges from 1–9, averaging 1.8 years, with breeding performance peaking at 2.8 years (Bouwhuis et al. 2009; Woodman et al. 2022). Although there are some continuous changes with age (Bouwhuis et al. 2009), the main age effects on individual-level traits are captured by two age-classes: first-years (hereafter juveniles) and older (hereafter adults, Gosler 1993; Harvey et al. 1979; Perrins 1979). Great tits are annual breeders with a single breeding season April–June, when they generally undertake one breeding attempt (in some parts of their range second clutches can occur, Verhulst 1998; Visser et al. 2003). Data used here are from 32 populations (Figure 1), the geographical range of which represents a large part of the species’ breeding range (Sullivan et al. 2009). Generally, data collection at these sites included regular visits to nest-boxes during breeding to track reproductive attempts, individually mark chicks and adults, and record their morphometrics, sex and age. Age is based either on year of hatching for local birds, or plumage characteristics for immigrants, where juveniles and adults can be discriminated based on retention of juvenile feathers in the wing (Svensson 1992). Metadata for populations can be found through the Studies of Populations of Individual Birds (www.spibirds.org, Culina et al. 2021).

**Fig. 1.**
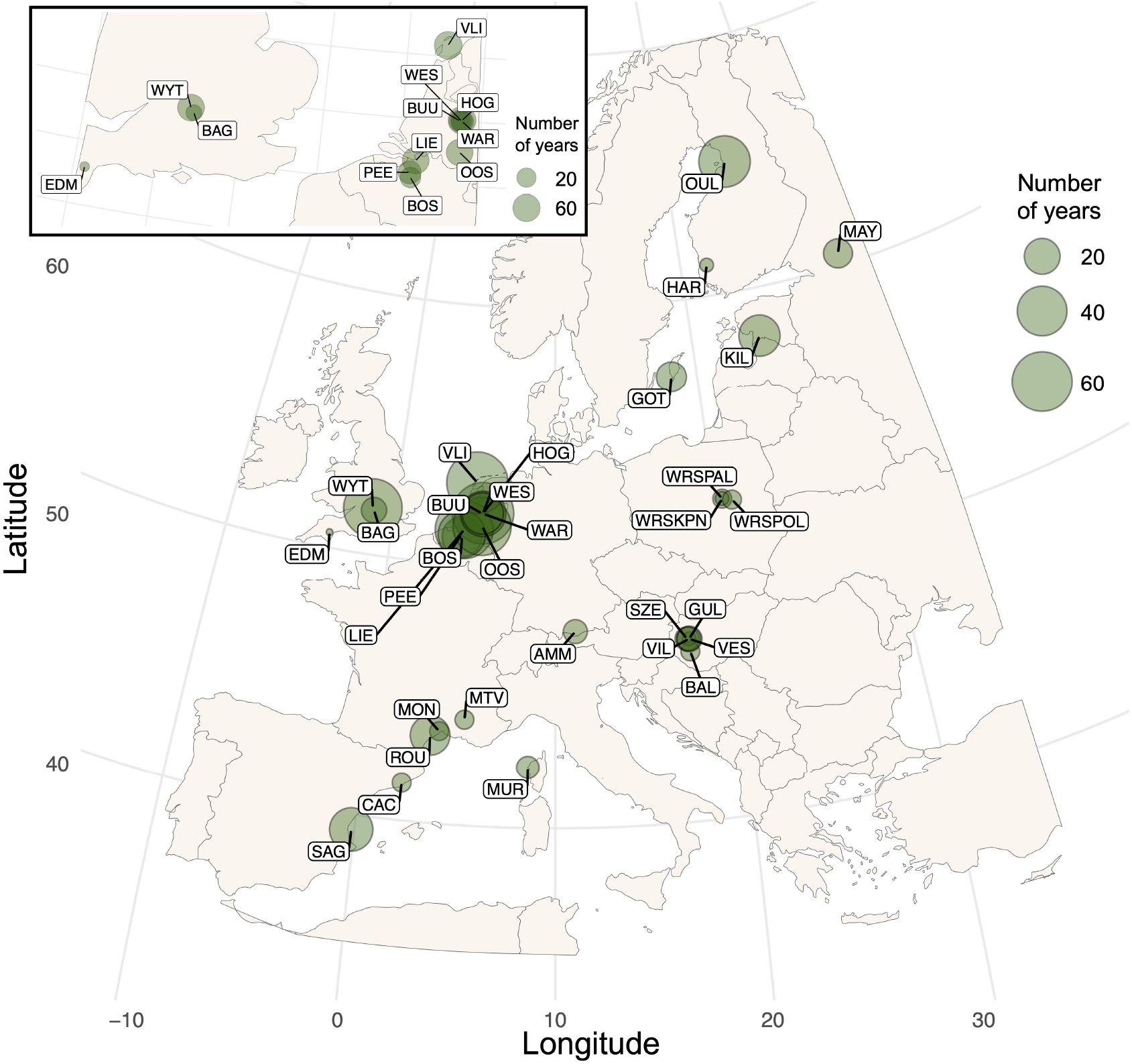
Map of the 32 great tit study populations across Europe, with point size relative to the number of years in the time series. The box in the top left shows the populations in The United Kingdom, Belgium, and The Netherlands in closer detail. Information for each study population can be found in Table S1.

### 2.2 Variation in age structure

We assigned age to all breeding great tits with known identity, across which exact year of hatching (birds caught within their first year) was known for 82.5% of 135,967 captures. Birds first captured with adult plumage were assigned a minimum age of 2 and included in analyses, with subsequent age estimates based on this (15.3% and 20.1% of females and males; given annual mortality rates *>*50% this assumption is likely to be accurate in most cases – also, in these cases, individuals can still accurately be designated as ‘juvenile’ or ‘adult’). Age was assigned to 62.1% of parents where at least one egg was laid (due to a combination of some studies’ protocols not always including adult identification, nests failing prior to adult capture, and unsuccessful trapping attempts, parental identity was unknown in some cases).

For each annual breeding population, we calculated the proportion of juveniles found breeding (additionally, we calculated: mean population age; proportion of senescent individuals; and change in these structures compared to a running mean as alternative descriptors of age structure, supporting information). We calculated this for every annual population, but only used data from years where the population included at least 20 individuals (mean, IQR: 230, 60–356) and *>*25% of the population was aged (mean, IQR: 56.0%, 36.1–78.2%). In total, the study period spanned 1956–2022, collectively consisting of 702 study years and 131,150 captures of 77,964 breeding individuals.

### 2.3 Reproductive and environmental variables

We assessed how reproductive and environmental variables that vary at different spatial scales relate to variation in age structure. First, we considered the influence of within-population average clutch size in year *t* − 1 on population-specific age structure in year *t*. We would expect variation in annual mean clutch size to affect the age structure of the following breeding season, where large average clutch sizes would lead to more recruits (Ahola et al. 2009) and therefore a higher proportion of breeding juveniles in the next year, thus we test this prediction here. We calculated population-level average clutch size as the mean number of eggs produced per breeding attempt within a breeding season.

Second, we considered two climatic variables: temperature and precipitation. We calculated the average mean daily temperature (°C) and the average daily precipitation sum (mm) across four periods preceding the focal breeding season for each population: June–August (hereafter summer); September–November (autumn); December–February (winter); and March–May (spring). We also considered the frequency of extreme climatic events (ECEs) by calculating the number of ‘cold ECEs’ June–May; and the number of ‘hot ECEs’ June–May. We define ECEs as events with a *<*5% probability of occurrence across the entire study period (1956–2022) in each population separately (Marrot et al. 2017; Moreno & Møller 2011). Thus, a cold ECE occurred when minimum daily temperature reaches below the 5% threshold June–May; and a hot ECE as when maximum daily temperature reaches above the 95% threshold.

Third, we considered European beech *Fagus sylvatica* masting, an environmental variable which is generally understood to vary at a large spatial scale. Beech masting is the annual production of seeds (Kelly 1994), which constitute part of the winter diet of great tits, thus influencing survival, particularly among juveniles (Källander 1981; Perdeck et al. 2000). The distribution of beech does not extend across the entire range of populations assessed here, and in southern Europe is restricted to higher altitudes (Bolte et al. 2007). However, masting-related demographic fluctuations in tits are synchronised across regions with and without beech. This suggests beech masting may be correlated with fruiting of other tree species important for survival, such that years with a large beech crop are rich in other food resources, promoting survival across different habitats (Klomp 1980; Perrins 1966). Thus, for each annual population, we obtained a masting value from a long-term continental-scale dataset of beech masting time series up to 2017 (MASTREE+, Hacket-Pain et al. 2022), using the masting value from a data collection site closest to that of each population in the year preceding breeding. The central coordinates for all sites were less than 1500km from the focal breeding population, which is the spatial scale at which masting remains synchronised (Bogdziewicz et al. 2021), and most were much closer (median, IQR: 143.2km, 87.6–297.0km). To assess the influence of masting at a more local scale, we created a subset of populations within the distribution of beech (Figure S1) and where annual data was collected within 100km of the population (12 populations, n = 188 population-years; further details for reproductive and environmental variables in supporting information).

### 2.4 Temporal variation in age structure and explanatory variables

First, we investigated the effects of the reproductive and environmental variables on population age structure. For each explanatory variable we constructed a linear mixed-effects model of the form

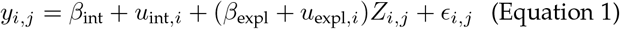

where *y* is the proportion of juveniles per population *i* and year *j, β*_int_ is an intercept, *u*_int,*i*_ denotes random intercepts for each population assumed to have a normal prior distribution with mean 0 and standard deviation 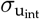, *β*_expl_ is a slope for the explanatory variable, *u*_expl,*i*_ denotes random slopes for the explanatory variable for each population also assumed to have a normal prior distribution, *Z*_*i*,*j*_ is the explanatory variable for each annual population, and *ϵ*_*i*,*j*_ is the residual error, assumed to have a normal prior distribution. This model was run for the 13 explanatory variables separately, as many of the environmental variables are highly correlated, thus leading to multicollinearity issues and making the interpretation of individual effects challenging.

These models were run using *brms* version 2.18.0 (Bü rkner 2017). We used default priors and ran four Markov chains for 6000 iterations with a burn-in of 3000, resulting in 12000 posterior samples. Chain convergence was evaluated using the diagnostic 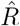 and effective sample size (Vehtari et al. 2021). We also ran the same models using alternative age structure descriptors (supporting information). The explanatory variables and age structure descriptors were z-normalised such that their relative effects could be assessed.

### 2.5 Spatial synchrony of variation in age structure

Second, we analysed whether age structure fluctuations are spatially synchronous, and whether this is explained by variation in the reproductive and environmental variables. Following Engen et al. (2005), we calculated a spatial auto-correlation function of the form

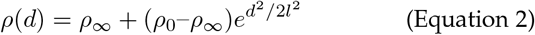

where *ρ*_0_ and *ρ*_*∞*_ are estimates of the correlation of age structure as distance approaches zero and infinity, respectively; 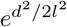 is a Gaussian positive definite autocorrelation function where *d* is distance between populations (in kilometres), and *l* is the standard deviation representing a standardised measure of the scale of spatial autocorrelation, i.e. the characteristic spatial scale at which temporal variation of an ecological property remains correlated (Jarillo et al. 2018; Lande et al. 1999). This model assumes that the spatial autocorrelation structure is Gaussian and that the parameters *ρ*_0_, *ρ*_*∞*_ and *l* are positive (as described in Engen et al. 2002; Lande et al. 1999). While it is possible that correlation in age structure between any two populations is negative, here we assume that correlation on average cannot be below zero. Modelling negative correlations on average is possible through non-parametric approaches (e.g. Bjørnstad et al. 1999b), but it is well established that using the parametric approach applied here is beneficial when assessing widescale spatial synchrony in ecological variables (Bjørnstad et al. 1999a; Engen et al. 2005; Vriend et al. 2023). The advantage is that that the model parameters have a biological interpretation which can be compared when considering synchrony of various demographic traits and, importantly, potential drivers of this synchrony can be quantitatively assessed.

The normalised age structure variables of all populations in each year were assumed to have a multivariate normal distribution where 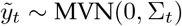. The diagonal elements of the variance-covariance matrix were set to 1, and the off-diagonal elements were defined by *ρ*_0_, *ρ*_*∞*_ and *l* given distance *d* between populations. Data from different populations were collected over a variable number of years, thus time series that overlap for longer were given more weight in the likelihood calculation. Overall log-likelihood was the sum of annual log-likelihoods optimised numerically to provide estimates for *ρ*_0_, *ρ*_*∞*_ and *l*. The distributions of these spatial synchrony parameters were obtained by parametric bootstrapping involving simulation from the multivariate normal distribution, based on the yearly set of populations in the data and the estimated spatial synchrony parameters (Engen et al. 2005; Lillegård et al. 2005). This was undertaken 2000 times, resulting in 2000 bootstrap replicates. The multi-variate normal distribution was constructed using *mvtnorm* version 1.1-3 (Genz et al. 2021).

Finally, we assessed the influence of reproductive and environmental variables in explaining spatial synchrony of age structure. Following previous methods (e.g. Grøtan et al. 2005; Sæther et al. 2007; Vriend et al. 2023), the proportion of juveniles in each annual population was regressed against population-specific explanatory variables in separate linear models. The residuals from these were then normalised and used as the variable of interest in the spatial synchrony model (Equation 2). This allowed us to calculate synchrony in age structure once the effects of the explanatory variables have been accounted for. All analysis was run in R statistical software version 4.2.2 (R Core Team 2021).

## 3 Results

### 3.1 Temporal variation in age structure and explanatory variables

We found marked temporal variation in age structure, with the annual proportion of juveniles ranging 0–0.89 across the 32 breeding populations over 1956–2022, and a very weak trend toward smaller proportions of juveniles found breeding over time (n = 702 population-years; supporting information; Figure S3a). We found that increased average annual clutch sizes were associated with a larger proportion of breeding juveniles the following year (posterior mode [95% credible intervals]: 0.437 [0.310, 0.553]).

Variation in age structure related to climatic factors, where breeding populations with a smaller proportion of juveniles followed warmer summers (-0.216 [-0.405, -0.038]) and winters (-0.250 [-0.436, -0.053]), and years with more frequent hot ECEs (-0.101 [-0.186, -0.016]). However, we found no effect of precipitation. Breeding populations had higher proportions of juveniles in years that followed winters with a large beech crop (0.225 [0.130, 0.314]). Moreover, this positive relationship was stronger when only assessing populations within 100km of the beech data collection site (0.365 [0.204, 0.538]; Figure 2; Table S2).

**Fig. 2.**
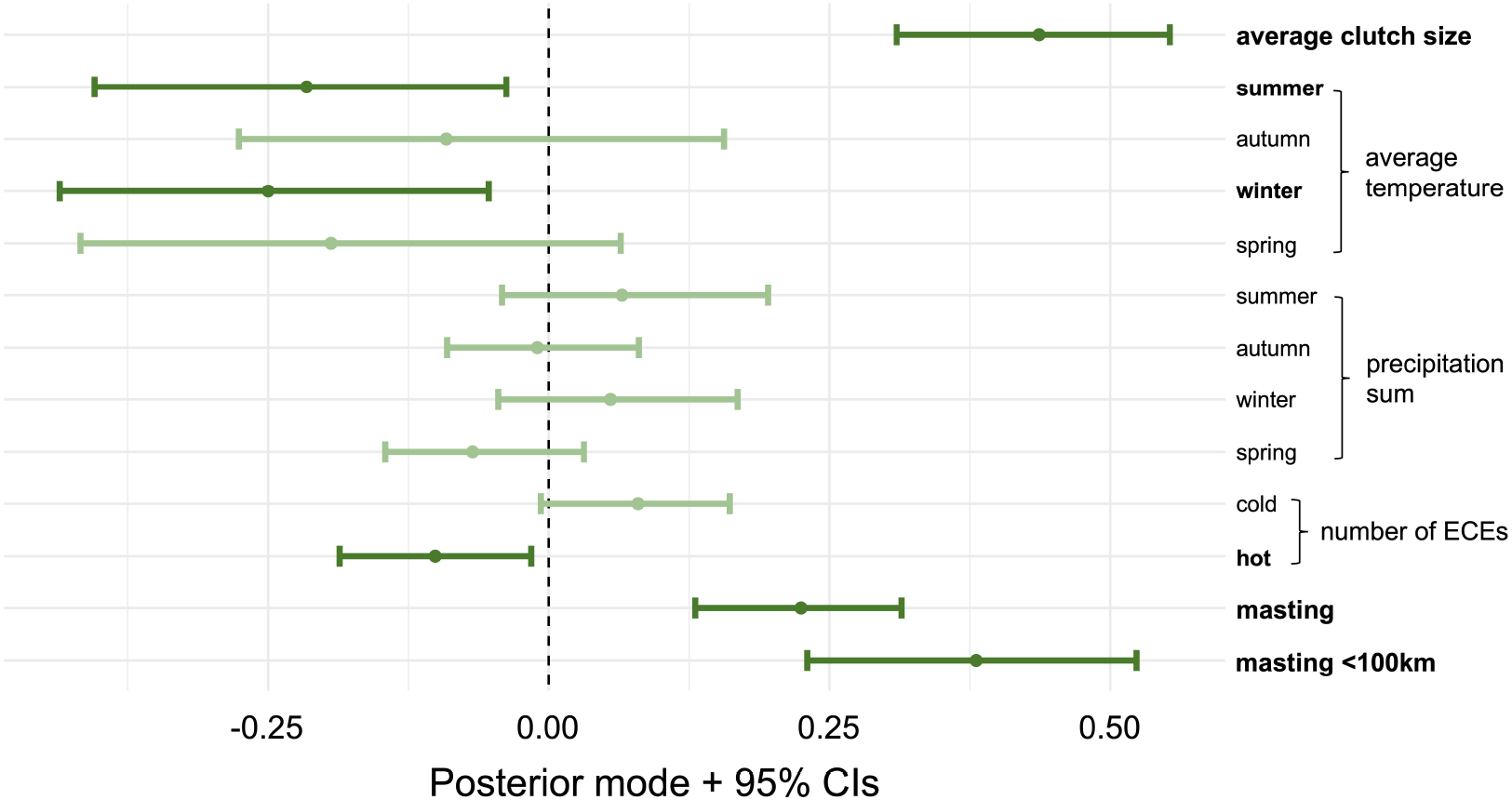
Posterior modes obtained from linear mixed-effects models which analyse the association between temporal variation in age structure and 13 reproductive and environmental variables across 32 great tit populations. Each point corresponds to a specific predictor variable (on the y-axis), and error bars denote 95% credible intervals. Points and error bars are reduced in saturation when credible intervals overlap zero, and explanatory variable text is bolded when they do not.

### 3.2 Spatial synchrony of variation in age structure

We found moderate spatial synchrony in age structure, with synchrony decreasing as distance between populations increased (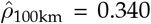 [0.260, 0.416]; 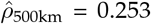 [0.163,0.330]; 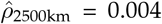 [*<*0.001, 0.112]; Figure 3a; Table 1), and the estimate for the scale of spatial autocorrelation was relatively large (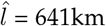[371km, 1000km]).

**TABLE 1.** Spatial synchrony of temporal variation in age structure across great tit breeding populations. Estimates are provided for spatial synchrony parameters (calculated in Equation 2); and for synchrony at distances of 100km, 500km, 1000km and 2500km

| Parameter | Median | 95% credible intervals |
| --- | --- | --- |
| $\rho_0$ | 0.344 | [0.264, 0.424] |
| $\rho_\infty$ | <0.001 | [<0.001, 0.112] |
| $l$ | 641km | [371km, 1000km] |
| $\rho_{100\text{km}}$ | 0.340 | [0.260, 0.416] |
| $\rho_{500\text{km}}$ | 0.253 | [0.163, 0.330] |
| $\rho_{1000\text{km}}$ | 0.115 | [0.035, 0.205] |
| $\rho_{2500\text{km}}$ | 0.004 | [<0.001, 0.112] |

**Fig. 3.**
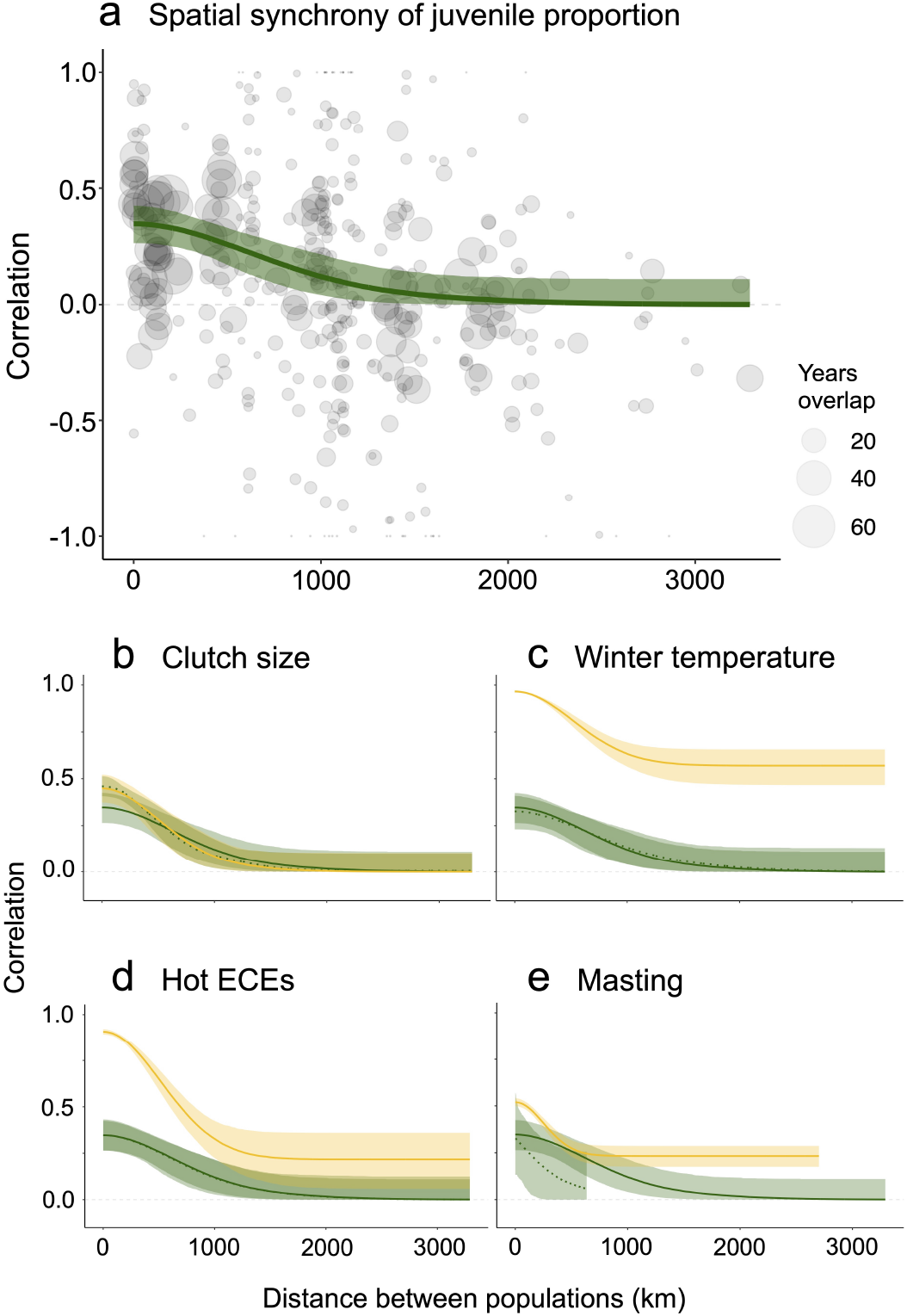
Spatial synchrony of temporal variation in age structure across great tit breeding populations in relation to distance between populations. In all plots, distance between populations (km) is on the x-axis and correlation between paired sites is on the y-axis. (a) Shows spatial synchrony of temporal fluctuations in the proportion of juveniles, where the green line is the median estimate of spatial synchrony (calculated in Equation 2) based on 2000 bootstrap replicates, with light green shading representing 95% credible intervals, and point size relative to the number of years of overlap between time series of pairwise sites. In (b–e), the dark green solid line is the estimate of spatial synchrony in the proportion of juveniles with 95% credible intervals, the yellow line is the spatial synchrony of the given reproductive or environmental variable, and the green dashed line is the spatial synchrony in the proportion of juveniles once accounting for the given reproductive or environmental variable.

Interestingly, neither clutch size nor many of the environmental variables significantly explained spatial synchrony in age structure (Figure 3b–d; Table S3). There is some evidence that masting may have a synchronising effect on age structure fluctuations when only considering populations within 100km of mast data collection (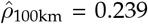 [0.030, 0.419];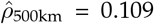 [*<*0.001, 0.303]; Figure 3e). However, these results should be interpreted with caution given that the synchrony estimates are based on a subset of populations across which the maximum distance between populations that overlap their time series (635km) is similar to the estimated spatial scale of synchrony in age structure across all populations.

## 4 Discussion

Using 131,150 captures of 77,964 individuals across 32 great tit populations collectively monitored over 702 years, we show that reproductive and environmental variables covary with breeding population age structure, with the strongest relationships between average clutch size from previous breeding, winter temperature, and variation in beech masting. We report moderate distant-dependent synchrony in age structure, with synchrony maintained at approximately 650km. However, despite association between reproductive and environmental factors with populations’ age structure, this does not explain overall spatial synchrony (except for some evidence that beech masting partially synchronises fluctuations in age structure).

### 4.1 Temporal variation in age structure and explanatory variables

Variation in age structure can have important consequences for demographic and social functioning (Coulson et al. 2005; Gamelon et al. 2019; Siracusa et al. 2023; Woodman et al. in press), yet there has been little research that links its variation across multiple populations directly to reproductive factors and environmental variability. This is because much research has focussed on how variation in population vital rates covary with environmental factors in single populations (Coulson et al. 2000, 2001; Farand et al. 2002), without explicitly linking this to variation in age structure (Hoy et al. 2020). Here, we provide evidence for an effect of reproductive and environmental variability in influencing breeding population age structure.

The strongest predictor of age structure was average clutch size, which predicts younger breeding populations the following year when clutch sizes are larger, suggesting that on average larger clutch sizes lead to more recruits (Ahola et al. 2009). Additionally, greater fecundity in great tits is linked to higher mortality (Payevsky 2006; Sæther 1988), thus there may be a relative increase in the proportion of juveniles the following year if there is high mortality among older individuals due to the cost of producing larger clutches. Directly linking variation in average clutch size and age structure reveals an important aspect of fluctuating population dynamics in great tits. This is because larger clutch sizes are produced when populations are smaller (Kluijver 1951; Lack 1952). Conversely, following an increase in the proportion of juveniles due to larger clutch sizes (as shown here), density-dependence will be strengthened, not only due to increasing population size, but also because juveniles have the strongest effect on density-dependent regulation, reducing recruitment and survival (Gamelon et al. 2016; Tinbergen et al. 1985) and producing fewer fledglings (Perrins & McCleery 1985).

However, perhaps the most interesting finding regarding the influence of average clutch size on age structure is that it has the greatest relative effect compared to environmental factors. This reveals that reproductive output is more influential for breeding population age structure than age-specific mortality induced by environmental variation. Of course, variation in clutch size itself is affected by environmental variability, such as temperature and food availability (Both & Visser 2005; Perrins 1965). However, in many species, age structure fluctuates considerably in response to environmental variation linked to mortality, because stressors such as extreme weather have more severe impacts on senescent and juvenile individuals compared to prime-aged individuals (Coulson et al. 2001; Farand et al. 2002; Hoy et al. 2015). Therefore, it is interesting that population-level reproductive output has a greater effect on age structure in this species compared to age-specific mortality events induced by environmental variability.

We report a relationship between age structure and local temperature, but not precipitation. Specifically, warmer summers and winters, and more frequent hot ECEs, are associated with a decrease in the proportion of juveniles. This is contrary to what might be expected where harsher winters would lead to elevated mortality of inexperienced juveniles. However, it has been shown that the reduction of fat following cold temperatures is not age-specific in great tits (Gosler 2002). Thus, mortality driven by colder winters might potentially even bias death in older individuals with lower basal metabolic rates (Broggi et al. 2010) or senescence in other physiological traits (Bouwhuis et al. 2012). A more plausible explanation is that cold temperature-driven mortality is non-age-specific, thus reducing local population size across all age-cohorts (van Balen 1980; Kluijver 1951; Payevsky 2006). In high-quality great tit habitats (as in many of the populations assessed here), there are often more individuals than available territories where more dominant individuals acquire the breeding sites (Perrins 1979). Great tits resident to a site are more dominant (Krebs 1982; Sandell & Smith 1991), thus upon their death, this likely makes more territories available to subdominant individuals migrating from surrounding lower-quality sites (Verhulst et al. 1997). This might therefore increase the number of immigrants found breeding following high rates of local mortality, for example, due to harsh winters (Grøtan et al. 2009; Tufto et al. 2005). Dispersal between birth and first breeding (natal dispersal) covers greater distances than dispersal between breeding attempts (Greenwood & Harvey 1982). Thus, great tits moving into new environments are often juveniles (Greenwood 1980). This therefore might generate an indirect relationship between colder winters and larger proportions of breeding juveniles the following spring. Further work should test this hypothesis by assessing winter temperature conditions under which immigration increases in great tits and other non-migratory species with sink populations in patchy environments.

We found larger beech mast events are followed by breeding seasons with a larger proportion of juveniles. Beech mast is important for winter survival in great tits, and previous work shows it particularly elevates juvenile survival (Clobert et al. 1988; Källander 1981; Perdeck et al. 2000). However, such studies assess either single populations, or populations close to one another, thus there is limited understanding of how important masting is for great tit demography on a continental-scale (but see Sæther et al. (2007) for its influence on size of populations located several hundred kilometres apart). Moreover, when we restricted analyses to populations that are within 100km of the beech data collection site, we found a greater effect of masting on breeding age structure. This could be due to two reasons: either data collected far from the focal population did not represent actual masting conditions experienced (although there is high spatial synchrony in beech crop cycles, Bogdziewicz et al. 2021); or such populations are in habitats with no or a lower density of beech, thus masting cannot influence demographic fluctuations. Indeed, when assessing population-specific trends, we find marked variation in the effect of masting on age structure (Figure S7l). Further, the strength of this relationship covaries with longitude, which approximates to the European distribution of beech (Figure S1; S8; S9). Although it has been suggested that masting-related tit population dynamics might be underpinned by synchronous fruiting of multiple tree species that elevate survival across wide-ranging habitats (Klomp 1980; Perrins 1966), our results may indicate that it is specifically beech which links variation in fruiting cycles with juvenile survival.

### 4.2 Spatial synchrony of variation in age structure

We found moderate spatial synchrony in breeding population age structure (Figure 3a). However, despite covariation between reproductive and environmental factors with age structure (Figure 2), accounting for such variables does not significantly affect the scale of spatial synchrony, other than evidence that beech masting may contribute to synchronising age structure fluctuations (Figure 3b–e; Table S3).

For any collection of populations, it is not necessarily expected that the same environmental variables would uniformly influence their age structures between years or synchronise fluctuations over time. An extensive body of demographic theory tells us that density-dependence can lead to complex dynamics, including chaos, in discrete population structures (Caswell 2000; Hastings et al. 1993; Levin 1981). Implicit in our analysis is the assumption that the populations have a fixed-point (stable) equilibrium age structure to which they gravitate, and that all populations are reasonably close to their equilibria. Even if the populations’ age structures do have a stable equilibrium, they are likely at different distances from their equilibria at any point in time, and thus going through different phases of transient dynamics (Hastings et al. 2018; Koons et al. 2005). If so, environmental effects may be obscured by the internal demographic processes that might be on divergent or uncorrelated trajectories. While these points serve as caveats concerning the complex dynamics influencing the populations, they also underscore the pronounced impact exerted by the explanatory variables (e.g. beech masting) on age structure, penetrating through the complicating dynamics.

Specifically, when only assessing populations within 100km of beech data collection sites, we find lower estimates of spatial synchrony in age structure once variation in masting has been accounted for. This may indicate that spatial synchrony in beech crop cycles (Bogdziewicz et al. 2021) act to synchronise fluctuations in age structure, but only in populations that breed within the distribution of beech. This is interesting given the effects of climate warming on beech masting, which has increased overall seed production, but reduced between-year variability and reproductive synchrony among individuals (Bogdziewicz et al. 2020). Thus, we might expect geographically distinct populations which are seed predators of beech, such as great tits, to have reduced synchrony in their age structure dynamics with increasing climate warming. This is particularly pertinent given that masting affects population dynamics across many taxa (Bogdziewicz et al. 2016).

Aside from this, it is striking that there is a lack of an effect from the other environmental variables in synchronising age structure. Spatial synchrony is generated either through dispersal between populations; interspecific trophic interactions with other spatially-synchronised populations; or a common influence from spatially autocorrelated environmental variables (the Moran effect). Dispersal between the assessed populations is unlikely to play a role here, as great tits disperse over smaller spatial scales compared to our estimate of the scale of synchrony in age structure (Greenwood et al. 1979; Tufto et al. 2005). However, dispersal more broadly might synchronise age structure fluctuations if there are simultaneous annual irruptive waves of young immigrants that move into the assessed sites prior to breeding (Nowakowski et al. 2003), especially seeing as annual variation in such waves correlate with years of high recruitment (Grøtan et al. 2009).

Our results may also suggest that the Moran effect does not underpin age structure spatial synchrony, because accounting for climatic variation does not reduce estimates of synchrony. This is pertinent because it suggests that although local environmental variability might induce changes to age structure within populations, broad-scale spatial autocorrelation in climate does not act to synchronise age structure between them. Thus, we would not expect widescale changes induced by climate change to synchronously affect age structure dynamics in this species.

Therefore, other factors may play a role in synchronising fluctuations in age structure. One factor potentially shaping this synchrony is spatial autocorrelation in ecological features that are related to, but not directly captured through the variation in, the environmental variables assessed here. For example, tits are highly susceptible to predation in the first month of fledging (Naef-Daenzer et al. 2001; Perrins & Geer 1980). Thus, if there is spatial synchrony in predator population dynamics, then this might induce synchrony in age structure across great tit populations through its effects on juvenile survival. Additionally, post-fledging food availability is particularly important for survival (Drent 1984; Payevsky 2006), which might also influence the proportion of juveniles found breeding the following season. Diet during this period consists predominantly of caterpillars (Verhulst & Hut 1996), the availability of which will not only be influenced by local weather, but other factors that affect invertebrate abundance and phenology, such as habitat heterogeneity and density-dependence. Exploring spatial synchrony in age structure while considering multiple trophic levels would be a particularly interesting perspective for future research.

Further, ecological features might not just act to synchronise population age structure through effects on age-specific survival, but also through their effects on reproduction. For example, great tit reproductive rates vary along an urban–natural gradient (Bukor et al. 2022; Charmantier et al. 2017), and reproductive responses to weather depend on whether breeding takes place in urban or natural habitats (Saulnier et al. 2023). Thus, if populations that are closer together experience more similar habitat types, this might induce spatial synchrony in their population dynamics.

Finally, we should expect that fluctuations in age structure are not fully explained by variation in environmental variables because of population-specific density-dependent dynamics. Local density-dependence can affect reproductive responses to environmental stochasticity (Møller et al. 2020), and differences in these dynamics between populations reduces spatial synchrony between them (Hugueny 2006; Walter et al. 2017). Thus, the spatial scale at which density-dependence remains similar between populations should influence the interaction between age structure and environmental variability, and the resultant spatial synchrony of fluctuations in age structure.

## 5 Conclusions

Using multiple overlapping time series from 32 wild great tit populations, our study quantifies associations between reproductive and environmental variables with age structure in spatially distinct breeding populations. Our findings emphasise the complex interaction between demographic processes and environmental variability, and we suggest that local ecological features and density-dependence will affect the dynamics of spatial synchrony between populations. Specifically, we report spatial synchrony of fluctuations in age structure at approximately 650km, but apart from some evidence of a synchronising effect of beech masting, we find that accounting for variation in reproductive and environmental factors does not reduce the scale of spatial synchrony. Further research should focus on how additional ecological features and population-specific density-dependent dynamics may contribute to the observed scale of spatial synchrony in age structure fluctuations that we find here.

## Acknowledgements

We are grateful to the hundreds of fieldworkers who have contributed to data collection in the 32 great tit populations assessed here over the past 67 years, without whom none of this work would be possible. We also thank the dedicated team at SPI-Birds, who continue to make strides towards open science allowing for cross-population studies such as this. Thank you also to Andrew Hacket Pain and Davide Ascoli et al., for their construction of the MASTREE+ dataset which allowed for our analysis of beech masting data, and also to Michał Bogdziewicz for his useful advice regarding beech synchrony. We acknowledge the E-OBS dataset from the EU-FP6 project UERRA (https://www.uerra.eu) and the Copernicus Climate Change Service, and the data providers in the ECA&D project (https://www.ecad.eu). JPW was supported by the Edward Grey Institute of Field Ornithology, University of Oxford. Additional funding acknowledged by co-authors is as follows: EA & EB from project PID2021-122171NB-I00/ AEI/10.13039/501100011033/ FEDER, UE; AA under state order to KarRC RAS No. FMEN-2022-0003; The Montpellier tit study (AC, SPC, AG & ML) has been funded since 2008 by the OSU-OREME and is part of the long-term Studies in Ecology and Evolution (SEE-Life) program of the CNRS; LC by the European Union’s Horizon 2020 research and innovation program under the Marie Skłodowska-Curie grant agreement No 838763; BD by the CNRS (PICS grants) and the ANR (ANR-06-JCJC-0082-01 and ANR-19-CE2-0007); TE from Research Council of Finland (SA338180); AL & GS from the HUN-REN TKI Hungarian Research Network, and by the Sustainable Development and Technologies National Programme of the Hungarian Academy of Sciences (FFT NP FTA); JCS from Ministry of Science and Innovation, Spain, project CGL-2020 PID2020-114907GB-C21; GS by the National Research, Development and Innovation Office (FK-137743) and the János Bolyai Research Scholarship of the Hungarian Academy of Sciences; JAF from BBSRC (BB/S009752/1), NERC (NE/S010335/1 and NE/V013483/1) and WildAI (CBR00730).

## References

Ahola, M.P., Laaksonen, T., Eeva, T. & Lehikoinen, E. (2009). Great tits lay increasingly smaller clutches than selected for: A study of climate- and density-related changes in reproductive traits. Journal of Animal Ecology, 78, 1298–1306.

van Balen, J.H. (1980). Population fluctuations of the Great Tit and feeding conditions in winter. Ardea, 55, 143–164.

Bjørnstad, O.N., Ims, R.A. & Lambin, X. (1999a). Spatial population dynamics: analyzing patterns and processes of population synchrony. Trends Ecol Evol, 14, 427–432.

Bjørnstad, O.N., Stenseth, N.C. & Saitoh, T. (1999b). Synchrony and scaling in dynamics of voles and mice in northern Japan. Ecology, 80, 622–637.

Bogdziewicz, M., Hacket-Pain, A., Ascoli, D. & Szymkowiak, J. (2021). Environmental variation drives continental-scale synchrony of European beech reproduction. Ecology, 102, 1–10.

Bogdziewicz, M., Kelly, D., Thomas, P.A., Lageard, J.G.A. & Hacket-Pain, A. (2020). Climate warming disrupts mast seeding and its fitness benefits in European beech. Nat Plants, 6, 88–94.

Bogdziewicz, M., Zwolak, R. & Crone, E.E. (2016). How do vertebrates respond to mast seeding? Oikos, 125, 300–307.

Bolte, A., Czajkowski, T. & Kompa, T. (2007). The north-eastern distribution range of European beech - a review. Forestry, 80, 413–429.

Both, C. & Visser, M.E. (2005). The effect of climate change on the correlation between avian life-history traits. Glob Chang Biol, 11, 1606–1613.

Bouwhuis, S., Choquet, R., Sheldon, B.C. & Verhulst, S. (2012). The forms and fitness cost of senescence: Age-specific recapture, survival, reproduction, and reproductive value in a wild bird population. American Naturalist, 179.

Bouwhuis, S., Sheldon, B.C., Verhulst, S. & Charmantier, A. (2009). Great tits growing old: Selective disappearance and the partitioning of senescence to stages within the breeding cycle. Proceedings of the Royal Society B: Biological Sciences, 276, 2769–2777.

Broggi, J., Hohtola, E., Koivula, K., Orell, M. & Nilsson, J. Å. (2010). Idle slow as you grow old: Longitudinal age-related metabolic decline in a wild passerine. Evol Ecol, 24, 177–184.

Bukor, B., Seress, G., Pipoly, I., Sándor, K., Sinkovics, C., Vincze, E., et al. (2022). Double-brooding and annual breeding success of great tits in urban and forest habitats. Curr Zool, 68, 517–525.

Bürkner, P.-C. (2017). brms: An R Package for Bayesian Multilevel Models Using Stan. J Stat Softw, 80, 1–28.

Caswell, Hal. (2000). Matrix population models: construction, analysis, and interpretation. 2nd ed. Sinauer Associates, Sunderland, Mass.

Charmantier, A., Demeyrier, V., Lambrechts, M., Perret, S. & Grégoire, A. (2017). Urbanization is associated with divergence in pace-of-life in great tits. Front Ecol Evol, 5.

Clobert, J., Perrins, C.M., McCleery, R.H. & Gosler, A.G. (1988). Survival Rate in the Great Tit Parus major in Relation to Sex, Age, and Immigration Status. Journal of Animal Ecology, 57, 287–306.

Coulson, T., Catchpole, E.A., Albon, S.D., Morgan, B.J.T., Pemberton, J.M., Clutton-Brock, T.H., et al. (2001). Age, sex, density, winter weather, and population crashes in Soay sheep. Science (1979), 292, 1528–1531.

Coulson, T., Gaillard, J.M. & Festa-Bianchet, M. (2005). Decomposing the variation in population growth into contributions from multiple demographic rates. Journal of Animal Ecology, 74, 789–801.

Coulson, T., Guinness, F., Pemberton, J. & Clutton-Brock, T.H. (2004). The demographic consequences of releasing a population of red deer from culling. Ecology, 85, 411–422.

Coulson, T., Milner-Gulland, E.J. & Clutton-Brock, T. (2000). The relative roles of density and climatic variation on population dynamics and fecundity rates in three contrasting ungulate species. Proceedings of the Royal Society B: Biological Sciences, 267, 1771–1779.

Culina, A., Adriaensen, F., Bailey, L.D., Burgess, M.D., Charmantier, A., Cole, E.F., et al. (2021). Connecting the data landscape of long-term ecological studies: The SPI-Birds data hub. Journal of Animal Ecology, 90, 2147–2160.

Drent, P.J. (1984). Mortality and dispersal in summer and its consequences for the density of Great Tits Parus major at the onset of autumn. Ardea, 72, 127–162.

Elton, C.S. (1924). Periodic Fluctuations in the Numbers of Animals: Their Causes and Effects. British Journal of Experimental Biology, 2, 119–163.

Engen, S., Lande, R. & Sæther, B.-E. (2002). The spatial scale of population fluctuations and quasi-extinction risk. Am Nat, 160, 439–451.

Engen, S., Lande, R., Seæther, B.-E. & Bregnballe, T. (2005). Estimating the pattern of synchrony in fluctuating populations. Journal of Animal Ecology, 74, 601–611.

Farand, É., Allainé, D. & Coulon, J. (2002). Variation in survival rates for the alpine marmot (Marmota marmota): Effects of sex, age, year, and climatic factors. Can J Zool, 80, 342–349.

Gamelon, M., Grøtan, V., Engen, S., Bjørkvoll, E., Visser, M.E. & Sæther, BE. (2016). Density dependence in an age-structured population of great tits: Identifying the critical age classes. Ecology, 97, 2479–2490.

Gamelon, M., Vriend, S.J.G., Engen, S., Adriaensen, F., Dhondt, A.A., Evans, S.R., et al. (2019). Accounting for interspecific competition and age structure in demographic analyses of density dependence improves predictions of fluctuations in population size. Ecol Lett, 22, 797–806.

Genz, A., Bretz, F., Miwa, T., Mi, X., Leisch, F., Scheipl, F., et al. (2021). Package ‘mvtnorm.’ Journal of Computational and Graphical Statistics, 11, 155.

Gosler, A.G. (2002). Strategy and constraint in the winter fattening response to temperature in the great tit Parus major. Journal of Animal Ecology, 71, 771–779.

Gosler, Andrew. (1993). The great tit. The great tit, Hamlyn species guides. Hamlyn, London.

Greenwood, P.J. (1980). Mating systems, philopatry and dispersal in birds and mammals. Anim Behav, 28, 1140–1162.

Greenwood, P.J. & Harvey, P.H. (1982). The natal and breeding dispersal of birds. Annu Rev Ecol Syst, 13, 1–21.

Greenwood, P.J., Harvey, P.H. & Perrins, C.M. (1979). The role of dispersal in the great tit (Parus major): The causes, consequences and heritability of natal dispersal. Journal of Animal Ecology, 48, 123–142.

Grøtan, V., Sæther, B., Engen, S., Balen, J.H. van, Perdeck, A.C. & Visser, M.E. (2009). Spatial and temporal variation in the relative contribution of density dependence, climate variation and migration to fluctuations in the size of great tit populations. Journal of Animal Ecology, 78, 447–459.

Grøtan, V., Sæther, B.-E., Engen, S., Solberg, E.J., Linnell, J.D.C., Andersen, R., et al. (2005). Climate causes large-scale spatial synchrony in population fluctuations of a temperate herbivore. Ecology, 86, 1472–1482.

Hacket-Pain, A., Foest, J.J., Pearse, I.S., LaMontagne, J.M., Koenig, W.D., Vacchiano, G., et al. (2022). MASTREE+: Time-series of plant reproductive effort from six continents. Glob Chang Biol, 28, 3066–3082.

Hansen, B.B., Grøtan, V., Herfindal, I. & Lee, A.M. (2020). The Moran effect revisited: spatial population synchrony under global warming. Ecography, 43, 1591–1602.

Harvey, P.H., Greenwood, P.J., Perrins, C.M. & Martin, A.R. (1979). Breeding success of great tits Parus major in relation to age of male and female parent. Ibis, 121, 216–219.

Hastings, A., Abbott, K.C., Cuddington, K., Francis, T., Gellner, G., Lai, Y.C., et al. (2018). Transient phenomena in ecology. Science (1979), 361.

Hastings, A., Hom, C.L., Ellner, S., Turchin, P. & Godfray, H.C.J. (1993). Chaos in ecology: is Mother Nature a strange attractor? Annu Rev Ecol Syst, 24, 1–33.

Heino, M., Kaitala, V., Ranta, E. & Lindstrom, J. (1997). Synchronous dynamics and rates of extinction in spatially structured populations. Proceedings of the Royal Society B: Biological Sciences, 264, 481–486.

Herfindal, I., Tveraa, T., Stien, A., Solberg, E.J. & Grøtan, V. (2020). When does weather synchronize life-history traits? Spatiotemporal patterns in juvenile body mass of two ungulates. Journal of Animal Ecology, 89, 1419–1432.

Hoy, S.R., MacNulty, D.R., Smith, D.W., Stahler, D.R., Lambin, X., Peterson, R.O., et al. (2020). Fluctuations in age structure and their variable influence on population growth. Funct Ecol, 34, 203–216.

Hoy, S.R., Petty, S.J., Millon, A., Whitfield, D.P., Marquiss, M., Davison, M., et al. (2015). Age and sex-selective predation moderate the overall impact of predators. Journal of Animal Ecology, 84, 692–701.

Hudson, P.J. & Cattadori, I.M. (1999). The Moran effect: a cause of population synchrony. Trends Ecol Evol, 14, 98–99.

Hugueny, B. (2006). Spatial synchrony in population fluctuations: Extending the Moran theorem to cope with spatially heterogeneous dynamics. Oikos, 115, 3–14.

Ims, R.A. & Andreassen, H.P. (2000). Spatial synchronization of vole population dynamics by predatory birds. Nature, 408, 194–196.

Jarillo, J., Sæther, B.E., Engen, S. & Cao, F.J. (2018). Spatial scales of population synchrony of two competing species: effects of harvesting and strength of competition. Oikos, 127, 1459–1470.

Jones, J., Doran, P.J. & Holmes, R.T. (2003). Climate and food synchronize regional forest bird abundances. Ecology, 84, 3024–3032.

Källander, H. (1981). The Effects of Provision of Food in Winter on a Population of the Great Tit Parus major and the Blue Tit P. caeruleus. Ornis Scandinavica, 12, 244–248.

Kelly, D. (1994). The evolutionary ecology of mast seeding. Trends Ecol Evol, 9, 465–470.

Kendall, B.E., Bjørnstad, O.N., Bascompte, J., Keitt, T.H. & Fagan, W.F. (2000). Dispersal, environmental correlation, and spatial synchrony in population dynamics. American Naturalist, 155, 628–636.

Klomp, H. (1980). Fluctuations and stability in great tit populations. Ardea, 68, 205–224.

Kluijver, H.N. (1951). The population ecology of the great tit, Parus m. major L. Ardea, 39, 1–135.

Koenig, W.D. (1999). Spatial autocorrelation of ecological phenomena. Trends Ecol Evol, 14, 22–26.

Koons, D.N., Grand, J.B., Zinner, B. & Rockwell, R.F. (2005). Transient population dynamics: Relations to life history and initial population state. Ecol Modell, 185, 283–297.

Koons, D.N., Iles, D.T., Schaub, M. & Caswell, H. (2016). A life-history perspective on the demographic drivers of structured population dynamics in changing environments. Ecol Lett, 19, 1023–1031.

Krebs, J.R. (1982). Territorial Defence in the Great Tit (Parus major): Do Residents Always Win? Behavioral Ecology and Sociobiology , 11, 185–194.

Lack, D. (1952). Reproductive Rate and Population Density in the Great Tit: Kluijvers Study. Ibis, 94, 167–173.

Lande, R., Engen, S. & Sæther, B.-E. (1999). Spatial Scale of Population Synchrony: Environmental Correlation versus Dispersal and Density Regulation. Am Nat, 154, 271–281.

Levin, S.A. (1981). Age-structure and stability in multiple-age spawning populations. In: Renewable Resource Management: Proceedings of a Workshop on Control Theory Applied to Renewable Resource Management and Ecology Held in Christchurch, New Zealand January 7–11, 1980. Springer, pp. 21–45.

Liebhold, A., Koenig, W.D. & Bjørnstad, O.N. (2004). Spatial synchrony in population dynamics. Annu Rev Ecol Evol Syst, 35, 467–490.

Lillegård, M., Engen, S. & Sæther, B.-E. (2005). Bootstrap methods for estimating spatial synchrony of fluctuating populations. Oikos, 109, 342–350.

Marrot, P., Garant, D. & Charmantier, A. (2017). Multiple extreme climatic events strengthen selection for earlier breeding in a wild passerine. Philosophical Transactions of the Royal Society B: Biological Sciences, 372.

Mills, L.. S. (2012). Conservation of wildlife populations: demography, genetics, and management. John Wiley & Sons.

Møller, A.P., Balbontín, J., Dhondt, A.A., Adriaensen, F., Artemyev, A., Bańbura, J., et al. (2020). Interaction of climate change with effects of conspecific and heterospecific density on reproduction. Oikos, 129, 1807–1819.

Moran, P.A.P. (1953). The statistical analysis of the Canadian Lynx cycle. II. Synchronization and Metereology. Aust J Zool, 1, 291–298.

Moreno, J. & Møller, A.P. (2011). Extreme climatic events in relation to global change and their impact on life histories. Curr Zool, 57, 375–389.

Mortelliti, A., Westgate, M., Stein, J., Wood, J. & Lindenmayer, D.B. (2015). Ecological and spatial drivers of population synchrony in bird assemblages. Basic Appl Ecol, 16, 269–278.

Naef-Daenzer, B., Widmer, F. & Nuber, M. (2001). Differential post-fledging survival of great and coal tits in relation to their condition and fledging date. Journal of Animal Ecology, 70, 730–738.

Nowakowski, J.K., Vähätalo, A. V. K N.J. & Vähätalo, A. (2003). Is the great tit Parus major an irruptive migrant in North-Eastern Europe? Ardea, 91, 231–243.

Olin, A.B., Banas, N.S., Wright, P.J., Heath, M.R. & Nager, R.G. (2020). Spatial synchrony of breeding success in the black-legged kittiwake Rissa tridactyla reflects the spatial dynamics of its sandeel prey. Mar Ecol Prog Ser, 638, 177–190.

Olmos, M., Payne, M.R., Nevoux, M., Prévost, E., Chaput, G., Du Pontavice, H., et al. (2020). Spatial synchrony in the response of a long range migratory species (Salmo salar) to climate change in the North Atlantic Ocean. Glob Chang Biol, 26, 1319–1337.

Ozgul, A., Childs, D.Z., Oli, M.K., Armitage, K.B., Blumstein, D.T., Olson, L.E., et al. (2010). Coupled dynamics of body mass and population growth in response to environmental change. Nature, 466, 482–485.

Paradis, E. (1997). Metapopulations and chaos: On the stabilizing influence of dispersal. J Theor Biol, 186, 261–266.

Paradis, E., Baillie, S.R., Sutherland, W.J. & Gregory, R.D. (1999). Dispersal and spatial scale affect synchrony in spatial population dynamics. Ecol Lett, 2, 114–120.

Paradis, E., Baillie, S.R., Sutherland, W.J. & Gregory, R.D. (2000). Spatial synchrony in populations of birds: Effects of habitat, population trend, and spatial scale. Ecology, 81, 2112–2125.

Parmesan, C. (2006). Ecological and Evolutionary Responses to Recent Climate Change. Annu Rev Ecol Evol Syst, 37, 637–669.

Payevsky, V.A. (2006). Mortality rate and population density regulation in the Great Tit, Parus major L.: A review. Russ J Ecol, 37, 180–187.

Perdeck, A.C., Visser, M.E. & Balen, J.H. van. (2000). Great Tit Parus major survival, and the beech-crop cycle. Ardea, 88, 99–106.

Perrins, C.M. (1965). Population fluctuations and clutch-size in the great tit, Parus major L. Journal of Animal Ecology, 34, 601–647.

Perrins, C.M. (1966). The effect of beech crops on great tit populations and movements. British Birds, 59, 419–432.

Perrins, C.M. (1979). British tits. 1st edn. Collins, London.

Perrins, C.M. & Geer, T.A. (1980). The effect of sparrowhawks on tit populations. Ardea, 68, 133–142.

Perrins, C.M. & McCleery, R.H. (1985). The effect of age and pair bond on the breeding success of Great Tits Parus major. Ibis, 127, 306–315.

R Core Team. (2021). R: a language and environment for statistical computing. Vienna: R Foundation for Statistical Computing.

Ranta, E., Kaitala, V., Lindstrõm, J. & Helle, E. (1997). The Moran Effect and Synchrony in Population Dynamics. Oikos, 78, 136–142.

Ripa, J. (2000). Analysing the Moran effect and dispersal: their significance and interaction in synchronous population dynamics. Oikos, 90, 175–187.

Rollinson, C.R., Finley, A.O., Alexander, M.R., Banerjee, S., Dixon Hamil, K.A., Koenig, L.E., et al. (2021). Working across space and time: non-stationarity in ecological research and application. Front Ecol Environ, 19, 66–72.

Ruxton, G.D. (1994). Low Levels of Immigration between Chaotic Populations can Reduce System Extinctions by Inducing Asynchronous Regular Cycles. Proc Biol Sci, 256, 189–193.

Sæther, B.-E. (1988). Pattern of covariation between life-history traits of European birds. Nature, 331, 616–617.

Sæther, B.-E., Engen, S., Grøtan, V., Fiedler, W., Matthysen, E., Visser, M.E., et al. (2007). The extended Moran effect and large-scale synchronous fluctuations in the size of great tit and blue tit populations. Journal of Animal Ecology, 76, 315–325.

Sandell, M. & Smith, H.G. (1991). Dominance, prior occupancy, and winter residency in the great tit (Parus major). Behav Ecol Sociobiol, 29, 147–152.

Saulnier, A., Bleu, J., Boos, A., Millet, M., Zahn, S., Ronot, P., et al. (2023). Reproductive differences between urban and forest birds across the years: importance of environmental and weather parameters. Urban Ecosyst, 26, 395–410.

Selås, V. (1997). Cyclic Population Fluctuations of Herbivores as an Effect of Cyclic Seed Cropping of Plants: The Mast Depression Hypothesis. Oikos, 80, 257–268.

Sibly, R.M. & Hone, J. (2002). Population growth rate and its determinants: An overview. Philosophical Transactions of the Royal Society B: Biological Sciences, 357, 1153–1170.

Siracusa, E.R., Pereira, A.S., Brask, J.B., Negron-Del Valle, J.E., Phillips, D., Platt, M.L., et al. (2023). Ageing in a collective: The impact of ageing individuals on social network structure. Philosophical Transactions of the Royal Society B: Biological Sciences, 378.

Sullivan, B.L., Wood, C.L., Iliff, M.J., Bonney, R.E., Fink, D. & Kelling, S. (2009). eBird: A citizen-based bird observation network in the biological sciences. Biol Conserv, 142, 2282–2292.

Svensson, L. (1992). Identification guide to European passerines. 4th, revd. edn. Lars Svensson; British Trust for Ornithology, Stockholm: Thetford.

Tinbergen, J.M., van Balen, J.H. & van Eck, H.M. (1985). Density dependent survival in an isolated Great Tit population: Kluyvers data reanalysed. Ardea, 73, 38–48.

Tufto, J., Ringsby, T.-H., Dhondt, A.A., Adriaensen, F. & Matthysen, E. (2005). A parametric model for estimation of dispersal patterns applied to five passerine spatially structured populations. Am Nat, 165, 13–26.

Vehtari, A., Gelman, A., Simpson, D., Carpenter, B. & Bürkner, P.-C. (2021). Rank-Normalization, Folding, and Localization: An Improved 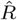 for Assessing Convergence of MCMC (with Discussion). Bayesian Anal, 16, 667–718.

Verhulst, S. (1998). Multiple breeding in the Great Tit, II. The costs of rearing a second clutch. Funct Ecol, 12, 132–140.

Verhulst, S. & Hut, R.A. (1996). Post-fledging care, multiple breeding and the costs of reproduction in the great tit. Animal Behavior, 51, 957–966.

Verhulst, S., Perrins, C.M. & Riddington, R. (1997). Natal dispersal of Great Tits in a patchy environment. Ecology, 78, 864–872.

Visser, M.E., Adriaensen, F., Van Balen, J.H., Blondel, J., Dhondt, A.A., Van Dongen, S., et al. (2003). Variable responses to large-scale climate change in European Parus populations. Proceedings of the Royal Society B: Biological Sciences, 270, 367–372.

Vriend, S.J.G., Grøtan, V., Gamelon, M., Adriaensen, F., Ahola, M.P., Álvarez, E., et al. (2023). Temperature synchronizes temporal variation in laying dates across European hole-nesting passerines. Ecology, 104.

Walter, J.A., Sheppard, L.W., Anderson, T.L., Kastens, J.H., Bjørnstad, O.N., Liebhold, A.M., et al. (2017). The geography of spatial synchrony. Ecol Lett, 20, 801–814.

Wan, X., Holyoak, M., Yan, C., Le Maho, Y., Dirzo, R., Krebs, C.J., et al. (2022). Broad-scale climate variation drives the dynamics of animal populations: a global multi-taxa analysis. Biological Reviews, 97, 2174–2194.

Woodman, J.P., Cole, E.F., Firth, J.A., Perrins, C.M. & Sheldon, B.C. (2022). Disentangling the causes of age-assortative mating in bird populations with contrasting life-history strategies. Journal of Animal Ecology.

Woodman, J.P., Gokcekus, S., Beck, K.B., Green, J.P., Nussey, D.H. & Firth, J.A. (2024). The ecology of ageing in wild societies: linking age structure and social behaviour. Philosophical Transactions of the Royal Society B: Biological Sciences, in press (accepted).

