## Supplementary material for "Continent-wide drivers of spatial synchrony in age structure across wild great tit populations": Figure S; supporting information; Table S

**Table of contents**


| Section | Page |
|---|---|
| 1. Supporting methods | 3 |
| Variation in age structure | 3 |
| Reproductive and environmental variables | 4 |
| 2. Supporting results | 6 |
| Temporal variation in age structure and explanatory variables | 6 |
| 3. Supplementary tables | 7 |
| 4. Supplementary figures | 16 |
| References | 28 |

### 1. Supporting methods

#### Variation in age structure

In addition to analysis on the proportion of breeding juveniles, we also calculated breeding population age structure through five alternative methods.

First, we calculated the mean breeding population age. As stated in the main text, exact year of hatching is known for all locally-hatched individuals that are ringed at the nest. Although annual immigration rates can be high in the assessed populations (mean, interquartile range: 69.7%, 57.2–88.3%), in many cases adults are first caught as juveniles (mean, IQR: 57.9%, 44.1–81.4%), and thus can be aged accurately. Therefore, exact age was known for 82.5% of 135,967 captures. For the remaining 17.5% of captured individuals (i.e. birds first caught with adult plumage), a minimum age of 2 was assigned, and subsequent age estimates were based on this (15.3% and 20.1% of females and males). This uncertainty does not affect accuracy of calculating our main age structure descriptor, because individuals can still be designated as ‘juvenile' or ‘adult’ with certainty. For the calculation of mean breeding population age, this assumption may affect accuracy of the age structure descriptor. However, given annual mortality rates >50% (Bouwhuis *et al.* 2009; Clobert *et al.* 1988), the assumption is likely to be accurate in the majority of cases. Given that individuals first encountered as adults in the first year of data collection in each population are all assigned an age of 2, the first three years of data collection were removed from each population in the analysis of mean breeding population age such that more accurate individual age estimates were used (n = 637, whereas n = 702 in the main analyses).

Second, we calculated the proportion of senescent individuals in the breeding population. We assume that senescence begins in great tits at 2.8 years (Bouwhuis *et al.* 2009); therefore, due to annual breeding, senescent individuals are defined as those of 3-years or older. Individuals assigned a minimum age of 2 in the first year of data collection which survive to the next year are accurately assigned as ‘senescent individual’ (because they must be 3-years or older). Thus, only the first year of data collection were removed from each population (n = 688).

Finally, we calculated $\delta_t$, the temporal deviations in the described structures (proportion of juveniles, mean breeding population age, and proportion of senescent individuals) compared to population-specific moving averages with a window size of 3 years. Specifically, $\delta_t = i_t - (\sum_{t-1}^{t-3} i_t)/3$, where $i_t$ is any of the above structural measures in year *t*. This metric addresses how the explanatory variables might induce momentary change in structure, beyond how those variables correlate with static measures of yearly structure. In order to compare the static age structure measure to a three-year average, data were restricted to populations with at least four years of continuous data. All age structure descriptors were highly correlated (Figure S2).

In addition to testing for associations between reproductive and environmental variables and variation in age structure, as in the main text, we also explored overall time trends in variation in population age structure. We did this by using the same linear mixed-effects model of the form

$$y_{i,j} = \beta_{\text{int}} + u_{\text{int},i} + (\beta_{\text{year}} + u_{\text{year},i})Z_{i,j} + \varepsilon_{i,j}$$

where $y$ is the age structure descriptor per population $i$ and year $j$, $\beta_{\text{int}}$ is an intercept, $u_{\text{int},i}$ denotes random intercepts for each population assumed to have a normal prior distribution with mean 0 and standard deviation $\sigma_{u_{\text{int}}}$, $\beta_{\text{year}}$ is a slope for the linear time trend, $u_{\text{year},i}$ denotes random slopes for the linear time trends of each population also assumed to have a normal prior distribution, $Z_{ij}$ is the time indicator per annual population, and $\varepsilon_{ij}$ is the residual error, assumed to have a normal prior distribution.

**Reproductive and environmental variables**

In the main text, we consider the role of reproductive and environmental variables that vary at different spatial scales on temporal variation in, and spatial synchrony of, population age structure. First, we considered the influence of average clutch size in year $t-1$ on age structure in year $t$. An individual's clutch size may vary in response to several demographic and environmental factors, such as local population size, predation risk, weather conditions, chick-rearing resource availability and parental quality (Boyce & Perrins 1987; Julliard *et al.* 1997; Møller *et al.* 2020; Perrins 1965; Pettifor *et al.* 2001). We would expect the variation in the within-year mean clutch size to affect the age structure of the following annual breeding population, where higher average clutch sizes would lead to greater numbers of recruits (Ahola *et al.* 2009) and therefore a higher proportion of breeding juveniles, thus we test this prediction here. Although the number of fledglings may provide a more accurate proxy for recruitment, data on fledgling number were less complete compared to clutch size across the populations, and the number of eggs laid strongly relates to the number of recruits at the pair-level (Perrins 1965; Perrins & Moss 1975).

We also consider the influence of climatic variables on breeding population age structure: temperature and precipitation, which have both been linked to great tit survival and reproduction (van Balen 1980; Bejer & Rudemo 1985; Bordjan & Tome 2014; Greño *et al.* 2007; Perrins 1965). We extracted daily temperature and precipitation records from a corresponding 0.1° × 0.1° grid cell in the E-OBS dataset version 27.0e for each population (Cornes *et al.* 2018). From these, we assessed the impact of weather on age structure by considering weather averages across given time periods (as outlined in the main text). We also considered the influence of the frequency of extreme climatic events (ECEs). Here, we define ECEs as an event with a 5% or less probability of occurrence across the entire study period (1956–2022) in each population separately (Bailey & van de Pol 2016; Marrot *et al.* 2017; Moreno & Møller 2011), which has been linked to great tit survival and reproduction (Regan & Sheldon 2023). Thus, a ‘cold ECE’ is defined as when the minimum daily temperature reaches below 5% threshold June–May; and a ‘hot ECE’ as when maximum daily temperature reaches above 95% threshold June–May.

We also considered an environmental variable which varies at a larger spatial scale by using masting data from European beech *Fagus sylvatica*. This variable represents the annual production of beech seeds (Kelly 1994; Silvertown 1980), where maximum production correlates negatively with summer temperatures two years previously, but positively with summer temperatures the year before (Vacchiano *et al.* 2017). Beech masting maintains significant synchrony at spatial scales up to 1500km (Bogdziewicz *et al.* 2021), and its variation has been linked to spatial synchrony of population size in tits (Sæther *et al.* 2007). We obtained data from a long-term continental-scale dataset of masting time series of beech up to 2017 (MASTREE+, Hacket-Pain et al., 2022). For each year of data collection for each breeding population, we extracted a masting value from the year prior to breeding from the site closest to that of the population. Beech masting was measured on an ordinal scale 1–5 where 1 represents lowest reproductive output (Hacket-Pain *et al.* 2022). The central coordinates for masting sites were all less than 1500km from the focal breeding population, which is the spatial scale at which beech masting remains synchronised (Bogdziewicz *et al.* 2021), and most were much closer (median, IQR: 143.2km, 87.6–297.0km). However, as set out in the main text, to assess the influence of beech masting measured at a more local spatial scale, we created a subset of the data including only annual populations where beech mast data was collected 100km or closer to the breeding site (13 populations, n = 223).

Finally, in addition to analysis outlined in the main text, we considered an environmental variable that varies at a continental scale by assessing the influence of the North Atlantic Oscillation (NAO) index. The NAO index is based on the difference in sea-level pressure between the subtropical high-pressure centre near the Azores and the subpolar low-pressure centre south and east of Greenland, the variation of which is linked to annual fluctuations in temperature and precipitation (Hurrell 1995; Lamb & Peppler 1987; Wanner *et al.* 2001). Generally, positive NAO values during the winter correspond to wetter, warmer weather and earlier springs in northern Europe and drier, warmer weather and advanced springs in southern Europe (Gordo & Sanz 2010; Post & Stenseth 1999). We extracted an annual winter NAO value from The Climate Data Guide (Hurrell & Phillips 2003; Schneider *et al.* 2013).

### 2. Supporting results

#### Temporal variation in age structure and explanatory variables

We found very weak but significant temporal trends in breeding population age structure. Specifically, we find that over time there was a slight decrease in the proportion of breeding juveniles (posterior mode [95% credible intervals]: -0.007 [-0.011, -0.002]) and a slight increase in the breeding population mean age (0.005 [<0.001, 0.010]). We found no significant time trend in the proportion of breeding senescent individuals (0.002 [-0.002, 0.007]); or on temporal deviations in the described structures (proportion of juveniles, mean breeding population age, and proportion of senescent individuals) compared to population-specific moving averages (>-0.001 [-0.003, 0.002]; <0.001 [-0.002, 0.003]; <0.001 [-0.001, 0.003], respectively).

We found similar associations between the reproductive and environmental variables with age structure whether defining age structure as the proportion of breeding juveniles (as in the main text; Figure 2), the mean breeding population age (Figure S4; Table S2), the proportion of senescent individuals (Figure S5; Table S2), or changes in these structures compared to a population-specific moving average (Figure S6; Table S2). In addition to the environmental variables assessed in the main text, we found that, slightly older breeding populations followed winters with higher NAO values (proportion of juveniles posterior mode [95% credible intervals]: -0.115 [-0.189, -0.028]). This may be interpreted similarly to our results found regarding increased winter temperatures, in that when winter NAO values are higher this is associated with warmer winters (see main text discussion).

### 3. Supplementary tables

Tables S1 – The 32 great tit populations used in this study, with information provided for: the location; the population identifier code (used in Figure 1 in the main text, in Figures S1 & S7, and in analysis); initials of data provider(s); latitude (Lat) and longitude (Lon), in decimal degrees; the time series of data used in this study (with total number of years in parentheses); the average within-year population size over the study period (estimated as the number of observed individuals + the number of inferred breeders, where it is assumed there are two individuals per breeding attempt); and the average within-year percentage of aged individuals (calculated as the number of aged individuals divided by the estimated breeding population size, which includes non-captured individuals). Metadata for each population can be found through the Studies of Populations of Individuals Birds (SPI-Birds; www.spibirds.org, Culina et al. 2021).

| Location | Population code | Data provider(s) | Lat | Lon | Time series (total years) | Mean pop. size | Mean % aged |
|---|---|---|---|---|---|---|---|
| Ammersee-Starnbergersee, Germany | AMM | ND | 47.5800 | 11.1400 | 2011-2019 (9) | 597 | 73.8 |
| Bagley Wood, United Kingdom | BAG | SRE | 51.7000 | -1.2500 | 2005-2014 (10) | 255 | 83.7 |
| Balatonfüred, Hungary | BAL | ALi, GS | 46.5700 | 17.5300 | 2014-2019 (6) | 47 | 69.5 |
| Boshoek, Belgium | BOS | FA, EM | 51.0800 | 4.3200 | 1994-2018 (25) | 465 | 74.2 |
| Buunderkamp, Netherlands | BUU | MEV | 52.0100 | 5.4500 | 1984-1992, 1995-2005, 2007-2014 (28) | 204 | 58.6 |
| Can Catà, Spain | CAC | JCS | 41.4600 | 2.1400 | 2003-2004, 2011, 2013, 2015-2016 (6) | 253 | 28.9 |
| East Dartmoor, United Kingdom | EDM | MDB | 50.5900 | -3.7200 | 2014-2015, 2018 (3) | 69 | 30.8 |
| Gotland, Sweden | GOT | LC, BD | 57.1000 | 18.2000 | 2005-2016, 2021-2022 (14) | 889 | 58.7 |
| Gulya-Domb, Hungary | GUL | ALi, GS | 47.0500 | 17.5300 | 2019-2022 (4) | 61 | 54.5 |
| Harjavalta, Finland | HAR | TE | 61.2000 | 22.1000 | 1991-1994 (4) | 343 | 33.9 |

| | | | | | | | |
|---|---|---|---|---|---|---|---|
| Hoge Veluwe, Netherlands | HOG | MEV | 52.0200 | 5.5100 | 1956-2018 (63) | 337 | 76.0 |
| Kilingi-Nõme, Estonia | KIL | ALe | 58.1500 | 24.9600 | 1972-1992, 1996-2001 (27) | 545 | 65.4 |
| Liesbos, Netherlands | LIE | MEV | 51.3500 | 4.4200 | 1956-1967, 1971-1974, 1976-2003, 2005-2011, 2014-2018 (56) | 116 | 63.8 |
| Mayachino, Russia | MAY | AA | 60.4600 | 32.4900 | 1983, 1985, 1990, 2001, 2007-2008, 2013-2016, 2018, 2020-2021 (13) | 29 | 70.4 |
| Montpellier City, France | MON | SPC, AC, AG, ML | 43.5900 | 3.8600 | 2013-2018 (6) | 275 | 50.8 |
| Mont Ventoux, France | MTV | SPC, AC, AG, ML | 44.1000 | 5.1600 | 1987-1991, 1993 (6) | 48 | 49.7 |
| Muro, France | MUR | SPC, AC, AG, ML | 42.3600 | 8.5800 | 2000-2005, 2008, 2012 (8) | 42 | 68.6 |
| Oosterhout, Netherlands | OOS | MEV | 51.5200 | 5.5000 | 1958, 1964-2018 (56) | 100 | 76.1 |
| Oulu, Finland | OUL | MO, SR, EV | 65.0500 | 25.5300 | 1970, 1972-1989, 1995, 1999-2021 (43) | 266 | 51.7 |
| Peerdsbos, Belgium | PEE | FA, EM | 51.1600 | 4.2900 | 1983-2007, 2009-2012, 2014-2018 (36) | 230 | 62.6 |
| Rouvière, France | ROU | SPC, AC, AG, ML | 43.4000 | 3.4000 | 1992-2006, 2008-2010, 2012-2018 (25) | 65 | 56.7 |
| Sagunto, Spain | SAG | EA, EB | 39.4200 | -0.1500 | 1993-2022 (30) | 184 | 86.0 |
| Szentgál, Hungary | SZE | ALi, GS | 47.0600 | 17.4100 | 2013-2021 (9) | 100 | 61.4 |
| Veszprém, Hungary | VES | ALi, GS | 47.0500 | 17.5400 | 2013-2022 (10) | 104 | 69.9 |

| | | | | | | | |
|---|---|---|---|---|---|---|---|
| Vilma-puszta, Hungary | VIL | ALi, GS | 47.0500 | 17.5200 | 2014-2022 (9) | 42 | 71.8 |
| Vlieland, Netherlands | VLI | MEV | 53.1700 | 5.0300 | 1956-2018 (63) | 349 | 79.2 |
| Warnsborn, Netherlands | WAR | KVO | 52.0000 | 5.5200 | 1983, 1987-2018 (33) | 91 | 71.9 |
| Westerheide, Netherlands | WES | KVO | 52.0100 | 5.5000 | 1992-2018 (27) | 359 | 67.1 |
| Warsaw Kampinos, Poland | WRSKPN | MS | 52.2123 | 20.4714 | 2018-2022 (6) | 39 | 62.1 |
| Warsaw Palmiry, Poland | WRSPAL | MS | 52.2211 | 20.4649 | 2018-2021 (4) | 31 | 63.3 |
| Warsaw Pole Mokotowskie, Poland | WRSPOL | MS | 52.1247 | 21.0698 | 2017-2022 (6) | 53 | 69.6 |
| Wytham Woods, United Kingdom | WYT | EFC, JAF, BCS, JPW | 51.7700 | -1.3400 | 1966-2022 (57) | 602 | 68.0 |

Table S2 – Associations between temporal variation in age structure and 14 reproductive and environmental variables across 32 great tit populations. Results are obtained from linear mixed-effects models of the form

$$y_{i,j} = \beta_{\text{int}} + u_{\text{int},i} + (\beta_{\text{expl}} + u_{\text{expl},i})Z_{i,j} + \varepsilon_{i,j} \text{ (Equation 1)}$$

where $y$ is the age structure descriptor per population $i$ and year $j$, $\beta_{\text{int}}$ is an intercept, $u_{\text{int},i}$ denotes random intercepts for each population assumed to have a normal prior distribution with mean 0 and standard deviation $\sigma_{u_{\text{int}}}$, $\beta_{\text{expl}}$ is a slope for the explanatory variable, $u_{\text{expl},i}$ denotes random slopes for the explanatory variable for each population also assumed to have a normal prior distribution, $Z_{ij}$ is the explanatory variable, and $\varepsilon_{ij}$ is the residual error, assumed to have a normal prior distribution. The posterior mode denotes the estimated effect size of the explanatory variable on the age structure descriptor ($\beta_{expl}$), drawn from 12000 posterior samples, with 95% credible intervals.

| Age structure descriptor | Variable | Posterior mode | 95% CrI |
|---|---|---|---|
| Proportion breeding juveniles | Clutch size | 0.437 | [0.310, 0.553] |
| | Summer temperature | -0.216 | [-0.405, -0.038] |
| | Autumn temperature | -0.091 | [-0.276, 0.156] |
| | Winter temperature | -0.250 | [-0.436, -0.053] |
| | Spring temperature | -0.194 | [-0.417, 0.064] |
| | Summer precipitation | 0.065 | [-0.042, 0.195] |
| | Autumn precipitation | -0.010 | [-0.091, 0.080] |
| | Winter precipitation | 0.055 | [-0.045, 0.168] |
| | Spring precipitation | -0.068 | [-0.146, 0.031] |
| | Cold ECEs | 0.080 | [-0.007, 0.161] |
| | Hot ECEs | -0.101 | [-0.186, -0.016] |
| | Masting | 0.225 | [0.130, 0.314] |
| | Masting <100km | 0.365 | [0.204, 0.538] |
| | NAO | -0.115 | [-0.189, -0.028] |
| Mean breeding population age | Clutch size | -0.450 | [-0.584, -0.323] |
| | Summer temperature | 0.275 | [0.068, 0.493] |
| | Autumn temperature | 0.216 | [-0.002, 0.418] |
| | Winter temperature | 0.180 | [-0.109, 0.414] |
| | Spring temperature | 0.227 | [-0.063, 0.500] |
| | Summer precipitation | -0.078 | [-0.214, 0.041] |
| | Autumn precipitation | 0.002 | [-0.085, 0.086] |
| | Winter precipitation | -0.009 | [-0.108, 0.093] |
| | Spring precipitation | 0.040 | [-0.047, 0.131] |
| | Cold ECEs | -0.060 | [-0.161, 0.051] |

| | | | |
|---|---|---|---|
| | Hot ECEs | 0.137 | [0.052, 0.230] |
| | Masting | -0.173 | [-0.250, -0.094] |
| | Masting <100km | -0.248 | [-0.519, -0.071] |
| | NAO | 0.094 | [0.009, 0.169] |
| Proportion senescent individuals | Clutch size | -0.390 | [-0.496, -0.289] |
| | Summer temperature | 0.173 | [-0.001, 0.350] |
| | Autumn temperature | 0.205 | [-0.011, 0.407] |
| | Winter temperature | 0.126 | [-0.146, 0.343] |
| | Spring temperature | 0.180 | [-0.082, 0.425] |
| | Summer precipitation | -0.026 | [-0.140, 0.088] |
| | Autumn precipitation | -0.005 | [-0.086, 0.076] |
| | Winter precipitation | -0.022 | [-0.121, 0.080] |
| | Spring precipitation | 0.031 | [-0.070, 0.122] |
| | Cold ECEs | -0.032 | [-0.133, 0.077] |
| | Hot ECEs | 0.098 | [0.005, 0.193] |
| | Masting | -0.081 | [-0.159, 0.002] |
| | Masting <100km | -0.117 | [-0.278, 0.046] |
| | NAO | 0.071 | [-0.009, 0.151] |
| Proportion juveniles temporal deviations | Clutch size | 0.190 | [0.086, 0.287] |
| | Summer temperature | -0.023 | [-0.139, 0.079] |
| | Autumn temperature | -0.009 | [-0.150, 0.109] |
| | Winter temperature | -0.091 | [-0.300, 0.020] |
| | Spring temperature | -0.035 | [-0.150, 0.073] |
| | Summer precipitation | 0.061 | [-0.034, 0.191] |
| | Autumn precipitation | -0.018 | [-0.107, 0.078] |
| | Winter precipitation | 0.014 | [-0.075, 0.119] |
| | Spring precipitation | -0.046 | [-0.137, 0.050] |
| | Cold ECEs | 0.056 | [-0.063, 0.150] |
| | Hot ECEs | -0.058 | [-0.156, 0.031] |
| | Masting | 0.284 | [0.162, 0.385] |
| | Masting <100km | 0.480 | [0.289, 0.766] |
| | NAO | -0.153 | [-0.252, -0.058] |
| Mean age temporal deviations | Clutch size | -0.215 | [-0.342, -0.101] |
| | Summer temperature | -0.002 | [-0.147, 0.235] |
| | Autumn temperature | 0.017 | [-0.107, 0.223] |
| | Winter temperature | 0.114 | [-0.017, 0.507] |
| | Spring temperature | 0.027 | [-0.098, 0.193] |
| | Summer precipitation | -0.060 | [-0.195, 0.049] |
| | Autumn precipitation | 0.007 | [-0.085, 0.116] |
| | Winter precipitation | 0.028 | [-0.097, 0.141] |
| | Spring precipitation | 0.074 | [-0.018, 0.182] |
| | Cold ECEs | -0.037 | [-0.169, 0.133] |
| | Hot ECEs | 0.064 | [-0.046, 0.194] |

| | | | |
|---|---|---|---|
| | Masting | -0.250 | [-0.362, -0.139] |
| | Masting <100km | -0.365 | [-0.590, -0.149] |
| | NAO | 0.109 | [0.008, 0.234] |
| Proportion senescents temporal deviations | Clutch size | -0.183 | [-0.358, -0.078] |
| | Summer temperature | 0.091 | [-0.035, 0.288] |
| | Autumn temperature | 0.036 | [-0.078, 0.276] |
| | Winter temperature | 0.060 | [-0.064, 0.251] |
| | Spring temperature | 0.060 | [-0.042, 0.231] |
| | Summer precipitation | -0.061 | [-0.184, 0.035] |
| | Autumn precipitation | 0.014 | [-0.078, 0.109] |
| | Winter precipitation | -0.055 | [-0.196, 0.058] |
| | Spring precipitation | 0.037 | [-0.061, 0.131] |
| | Cold ECEs | -0.005 | [-0.170, 0.231] |
| | Hot ECEs | 0.096 | [-0.002, 0.202] |
| | Masting | -0.096 | [-0.201, 0.016] |
| | Masting <100km | -0.203 | [-0.46, 0.000] |
| | NAO | 0.125 | [-0.027, 0.244] |

Table S3 – Spatial synchrony of temporal variation in the proportion of juveniles in great tit breeding populations after accounting for variation in reproductive and environmental variables. Estimates are provided for spatial synchrony parameters (calculated in Equation 2 in the main text); and for synchrony at distances of 100km, 500km, 1000km and 2500km.

| Variable accounted for | Parameter | Median | 95% CrI |
|---|---|---|---|
| Clutch size | $\rho_0$ | 0.302 | [0.215, 0.392] |
| | $\rho_\infty$ | <0.001 | [<0.001, 0.111] |
| | $l$ | 564km | [267km, 964km] |
| | $\rho_{100\mathrm{km}}$ | 0.297 | [0.212, 0.383] |
| | $\rho_{500\mathrm{km}}$ | 0.201 | [0.092, 0.282] |
| | $\rho_{1000\mathrm{km}}$ | 0.075 | [0.009, 0.175] |
| | $\rho_{2500\mathrm{km}}$ | 0.002 | [<0.001, 0.112] |
| Summer temperature | $\rho_0$ | 0.348 | [0.262, 0.433] |
| | $\rho_\infty$ | <0.001 | [<0.001, 0.127] |
| | $l$ | 618km | [314km, 1018km] |
| | $\rho_{100\mathrm{km}}$ | 0.343 | [0.258, 0.425] |
| | $\rho_{500\mathrm{km}}$ | 0.250 | [0.147, 0.330] |
| | $\rho_{1000\mathrm{km}}$ | 0.110 | [0.025, 0.215] |
| | $\rho_{2500\mathrm{km}}$ | 0.004 | [<0.001, 0.127] |
| Autumn temperature | $\rho_0$ | 0.339 | [0.251, 0.431] |
| | $\rho_\infty$ | <0.001 | [<0.001, 0.119] |
| | $l$ | 646km | [334km, 1046km] |
| | $\rho_{100\mathrm{km}}$ | 0.335 | [0.248, 0.423] |
| | $\rho_{500\mathrm{km}}$ | 0.251 | [0.153, 0.332] |
| | $\rho_{1000\mathrm{km}}$ | 0.118 | [0.032, 0.220] |
| | $\rho_{2500\mathrm{km}}$ | 0.006 | [<0.001, 0.119] |
| Winter temperature | $\rho_0$ | 0.326 | [0.231, 0.410] |
| | $\rho_\infty$ | <0.001 | [<0.001, 0.128] |
| | $l$ | 709 | [332km, 1199km] |
| | $\rho_{100\mathrm{km}}$ | 0.322 | [0.229, 0.402] |
| | $\rho_{500\mathrm{km}}$ | 0.251 | [0.149, 0.324] |
| | $\rho_{1000\mathrm{km}}$ | 0.133 | [0.040, 0.231] |
| | $\rho_{2500\mathrm{km}}$ | 0.010 | [<0.001, 0.130] |
| Spring temperature | $\rho_0$ | 0.344 | [0.262, 0.430] |
| | $\rho_\infty$ | <0.001 | [<0.001, 0.146] |
| | $l$ | 693km | [357km, 1107km] |
| | $\rho_{100\mathrm{km}}$ | 0.340 | [0.259, 0.424] |
| | $\rho_{500\mathrm{km}}$ | 0.265 | [0.168, 0.343] |
| | $\rho_{1000\mathrm{km}}$ | 0.139 | [0.043, 0.233] |
| | $\rho_{2500\mathrm{km}}$ | 0.010 | [<0.001, 0.147] |
| Summer precipitation | $\rho_0$ | 0.335 | [0.248, 0.421] |

| | | | |
|---|---|---|---|
| | $\rho_\infty$ | <0.001 | [<0.001, 0.126] |
| | $l$ | 672km | [339km, 1118km] |
| | $\rho_{100km}$ | 0.332 | [0.246, 0.414] |
| | $\rho_{500km}$ | 0.254 | [0.156, 0.332] |
| | $\rho_{1000km}$ | 0.127 | [0.033, 0.230] |
| | $\rho_{2500km}$ | 0.007 | [<0.001, 0.127] |
| Autumn precipitation | $\rho_0$ | 0.323 | [0.241, 0.412] |
| | $\rho_\infty$ | <0.001 | [<0.001, 0.119] |
| | $l$ | 612km | [310km, 1014km] |
| | $\rho_{100km}$ | 0.319 | [0.239, 0.403] |
| | $\rho_{500km}$ | 0.233 | [0.129, 0.314] |
| | $\rho_{1000km}$ | 0.107 | [0.022, 0.204] |
| | $\rho_{2500km}$ | 0.004 | [<0.001, 0.119] |
| Winter precipitation | $\rho_0$ | 0.347 | [0.258, 0.433] |
| | $\rho_\infty$ | <0.001 | [<0.001, 0.110] |
| | $l$ | 604km | [330km, 997km] |
| | $\rho_{100km}$ | 0.341 | [0.255, 0.426] |
| | $\rho_{500km}$ | 0.247 | [0.247, 0.328] |
| | $\rho_{1000km}$ | 0.104 | [0.104, 0.211] |
| | $\rho_{2500km}$ | 0.003 | [0.003, 0.111] |
| Spring precipitation | $\rho_0$ | 0.348 | [0.259, 0.432] |
| | $\rho_\infty$ | <0.001 | [<0.001, 0.133] |
| | $l$ | 662km | [338km, 1101km] |
| | $\rho_{100km}$ | 0.343 | [0.257, 0.426] |
| | $\rho_{500km}$ | 0.260 | [0.159, 0.335] |
| | $\rho_{1000km}$ | 0.127 | [0.035, 0.228] |
| | $\rho_{2500km}$ | 0.006 | [<0.001, 0.133] |
| Cold ECEs | $\rho_0$ | 0.327 | [0.242, 0.415] |
| | $\rho_\infty$ | <0.001 | [<0.001, 0.118] |
| | $l$ | 675km | [325km, 1148km] |
| | $\rho_{100km}$ | 0.323 | [0.239, 0.410] |
| | $\rho_{500km}$ | 0.247 | [0.146, 0.328] |
| | $\rho_{1000km}$ | 0.122 | [0.033, 0.220] |
| | $\rho_{2500km}$ | 0.007 | [<0.001, 0.120] |
| Hot ECEs | $\rho_0$ | 0.347 | [0.264, 0.432] |
| | $\rho_\infty$ | <0.001 | [<0.001, 0.125] |
| | $l$ | 632km | [343km, 1001km] |
| | $\rho_{100km}$ | 0.342 | [0.260, 0.425] |
| | $\rho_{500km}$ | 0.254 | [0.153, 0.334] |
| | $\rho_{1000km}$ | 0.117 | [0.028, 0.215] |
| | $\rho_{2500km}$ | 0.005 | [<0.001, 0.125] |
| Masting | $\rho_0$ | 0.272 | [0.185, 0.362] |
| | $\rho_\infty$ | <0.001 | [<0.001, 0.134] |
| | $l$ | 793km | [324km, 1440km] |

| | | | |
|---|---|---|---|
| | $\rho_{100km}$ | 0.269 | [0.183, 0.357] |
| | $\rho_{500km}$ | 0.221 | [0.132, 0.295] |
| | $\rho_{1000km}$ | 0.136 | [0.037, 0.220] |
| | $\rho_{2500km}$ | 0.019 | [<0.001, 0.137] |
| Masting < 100km | $\rho_0$ | 0.301 | [0.104, 0.573] |
| | $\rho_\infty$ | <0.001 | [<0.001, 0.261] |
| | $l$ | 255km | [19km, 437924km] |
| | $\rho_{100km}$ | 0.239 | [0.030, 0.419] |
| | $\rho_{500km}$ | 0.109 | [<0.001, 0.303] |
| | $\rho_{1000km}$ | NA | NA |
| | $\rho_{2500km}$ | NA | NA |
| NAO | $\rho_0$ | 0.315 | [0.228, 0.402] |
| | $\rho_\infty$ | <0.001 | [<0.001, 0.119] |
| | $l$ | 697km | [329km, 1220km] |
| | $\rho_{100km}$ | 0.311 | [0.226, 0.394] |
| | $\rho_{500km}$ | 0.240 | [0.140, 0.319] |
| | $\rho_{1000km}$ | 0.127 | [0.032, 0.227] |
| | $\rho_{2500km}$ | 0.009 | [<0.001, 0.120] |

### 4. Supplementary figures

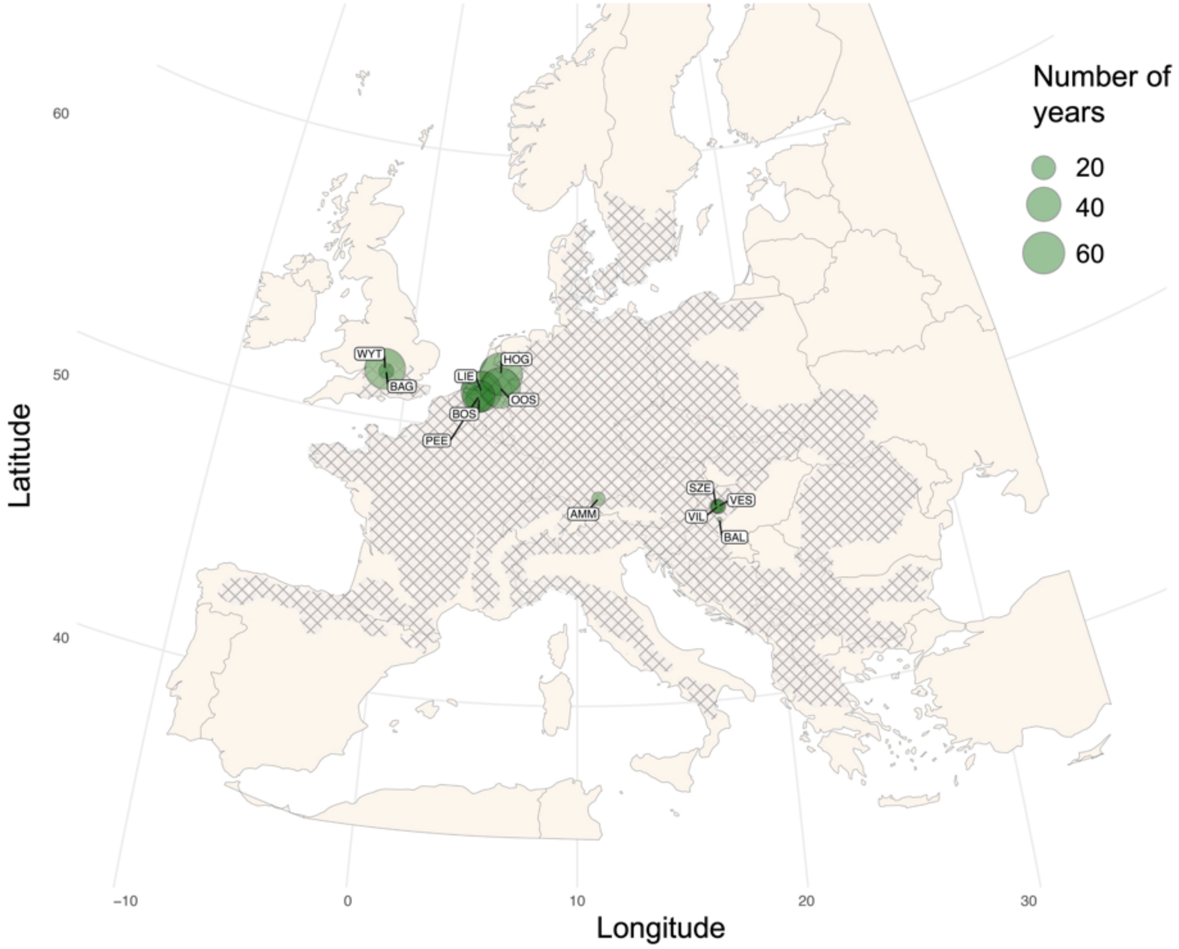


Figure S1 – Map of the 12 great tit study populations across Europe included in our sub-analysis assessing the influence of masting at a more local spatial scale. The annual populations include those that are within the continuous distribution range of beech (shown approximately in cross-hatching on the map, adapted from Bolte et al. 2007) and where the beech data was collected within 100km of the populations. Dark green points represent the great tit populations, with point size relative to the number of years in the time series.

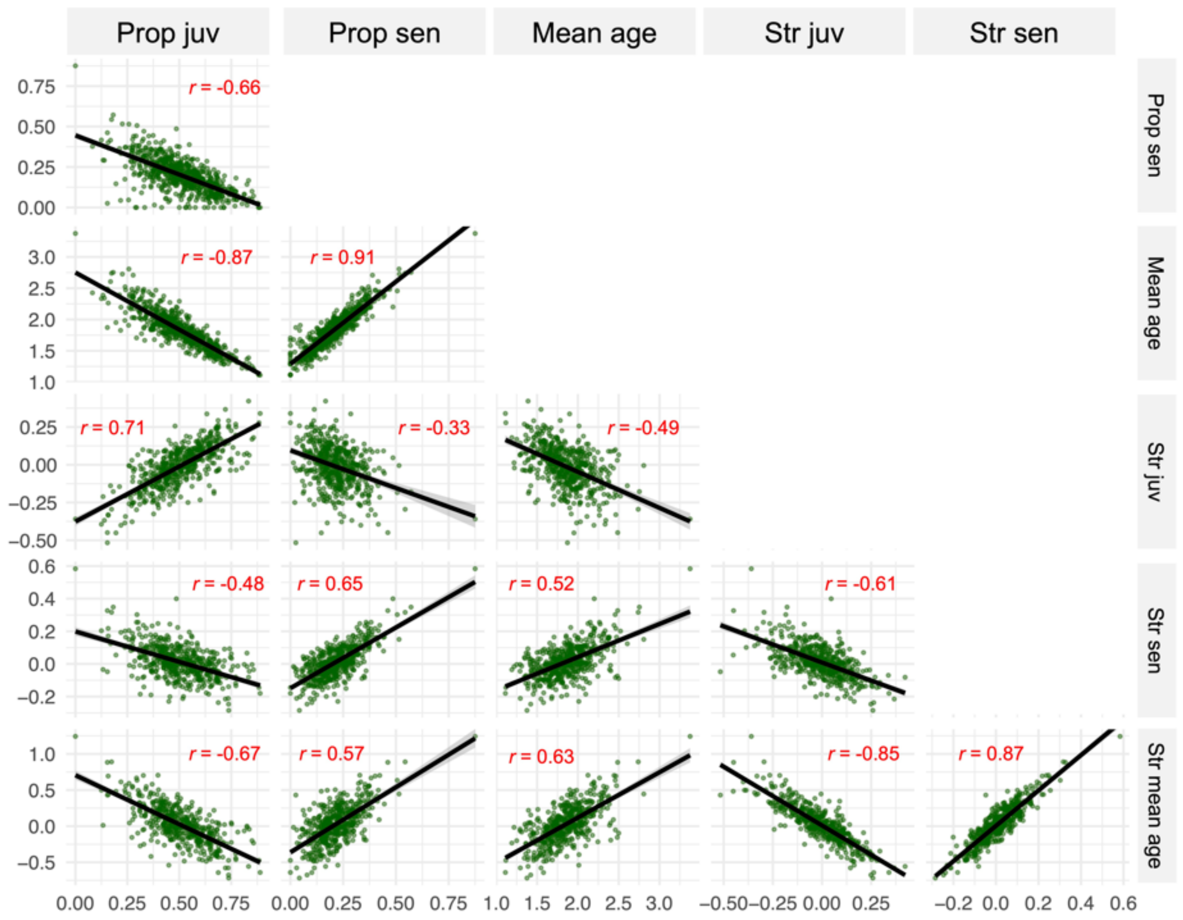


Figure S2 – Correlations between the different age structure descriptors (proportion of breeding juveniles, proportion of senescent individuals, mean breeding population age, and the temporal deviations in these three measures compared to population-specific moving averages) used in this study. In all plots, the black line is a linear regression which models the relationship between the two age structure measures, and the shading around this shows the 95% confidence intervals. The relationship is significant where $p < 0.05$ in all cases, and the $r$ value obtained from a Spearman's rank correlation labelled in red on all plots.

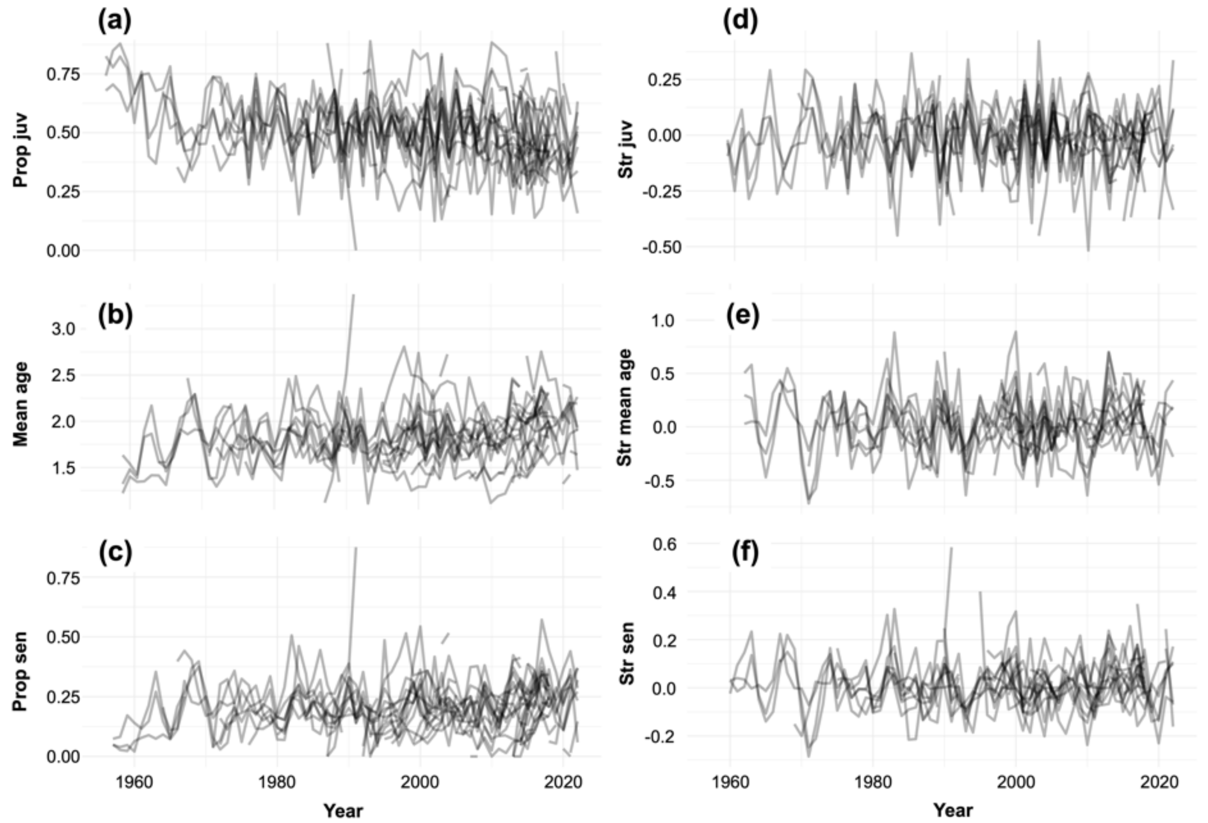


Figure S3 – Temporal variation in breeding population age structure across European great tit populations. In all plots and analysis, annual populations were only included if the population included at least 20 individuals (mean, IQR: 230, 60–356) and at least 25% of the population was aged (mean, IQR: 56.0%, 36.1–78.2%). Each line corresponds to a continuous time series from a single population (i.e. broken lines represent time series where some annual breeding populations consisted of either less than 20 individuals or less than 25% aged individuals). (a) Shows the proportion of breeding juveniles (32 populations, n = 702); (b) mean breeding population age (32 populations, n = 637); (c) proportion of breeding senescent individuals (32 populations, n = 688); (d) change in the proportion of juveniles compared to a 3-year running mean (30 populations, n = 549); (e) change in the mean breeding population age compared to a 3-year running mean (25 populations, n = 493); and (f) change in the proportion of senescent individuals compared to a running mean (27 populations, n = 536).

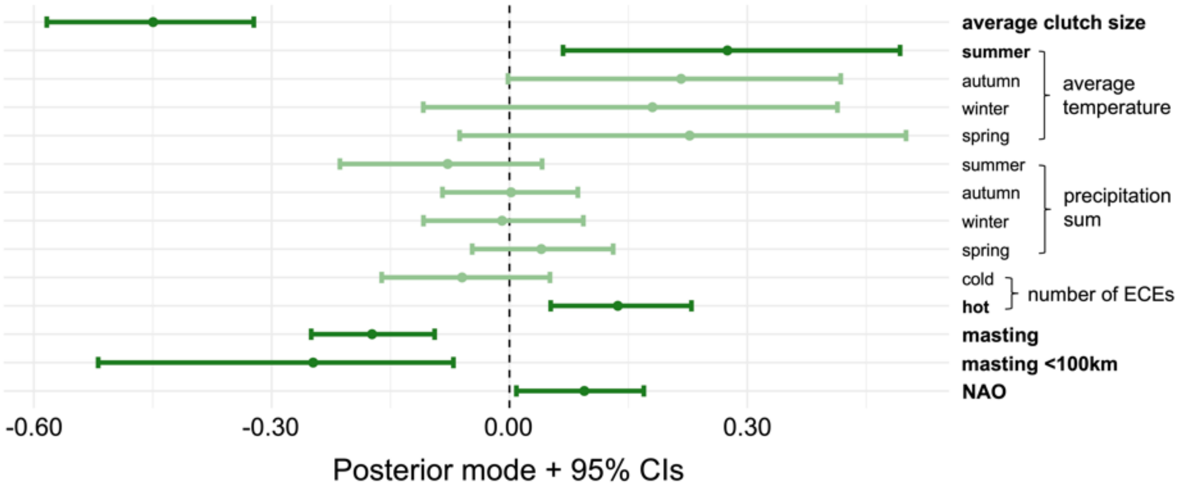


Figure S4 – Posterior modes obtained from linear mixed-effects models which analyse the association between temporal variation in mean breeding population age and 14 reproductive and environmental variables across 32 great tit populations. Each point corresponds to a specific predictor variable (on the y-axis), and error bars denote 95% credible intervals. Points and error bars are reduced in saturation when credible intervals overlap zero, and explanatory variable text is bolded when they do not.

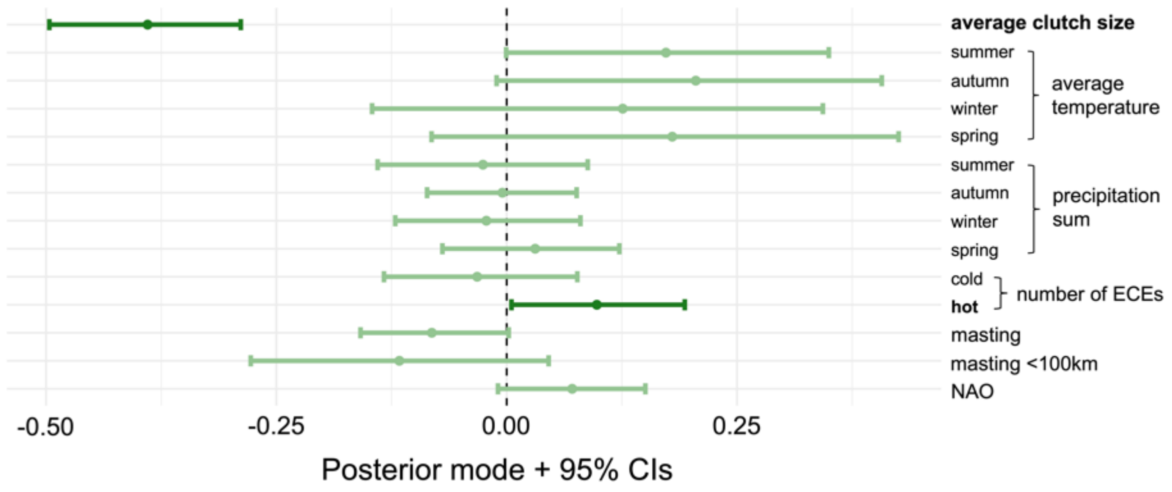


Figure S5 – Posterior modes obtained from linear mixed-effects models which analyse the association between temporal variation in the proportion of breeding senescent individuals and 14 reproductive and environmental variables across 32 great tit populations. Each point corresponds to a specific predictor variable (on the y-axis), and error bars denote 95% credible intervals. Points and error bars are reduced in saturation when credible intervals overlap zero, and explanatory variable text is bolded when they do not.

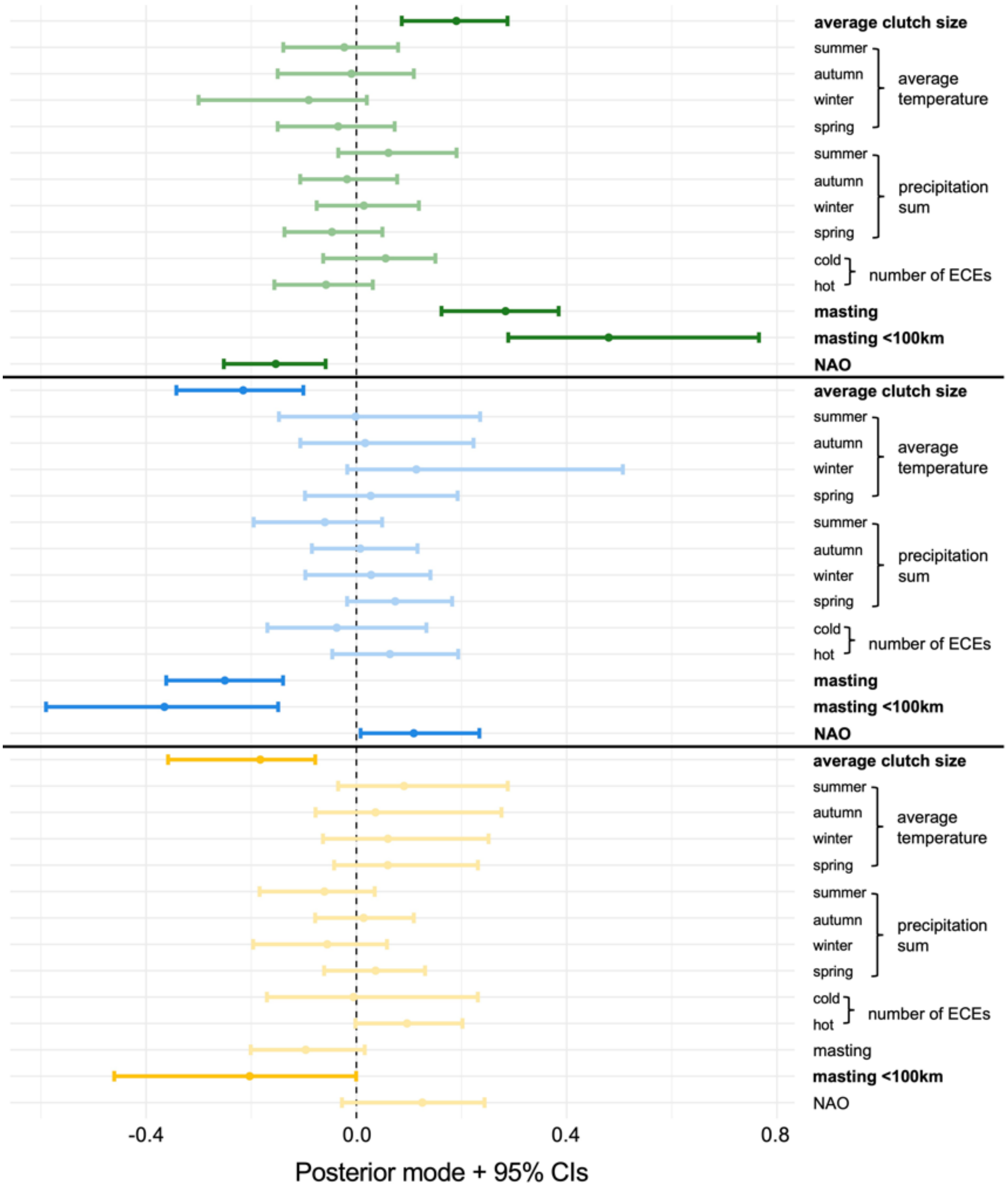


Figure S6 – Posterior modes obtained from linear mixed-effects models which analyse the association between temporal variation in breeding population age structure (defined as the difference between the within-year static age structure measure and that of a running average calculate as the mean in the three years previous) and 14 reproductive and environmental variables across 32 great tit populations. Each point corresponds to a specific predictor variable (on the y-axis), and error bars denote 95% credible intervals. Points and error bars are reduced in saturation when credible intervals overlap zero, and explanatory variable text is bolded when they do not. Green points are from analysis assessing the proportion of breeding juveniles, blue for the breeding population mean age, and yellow for the proportion of senescent individuals.

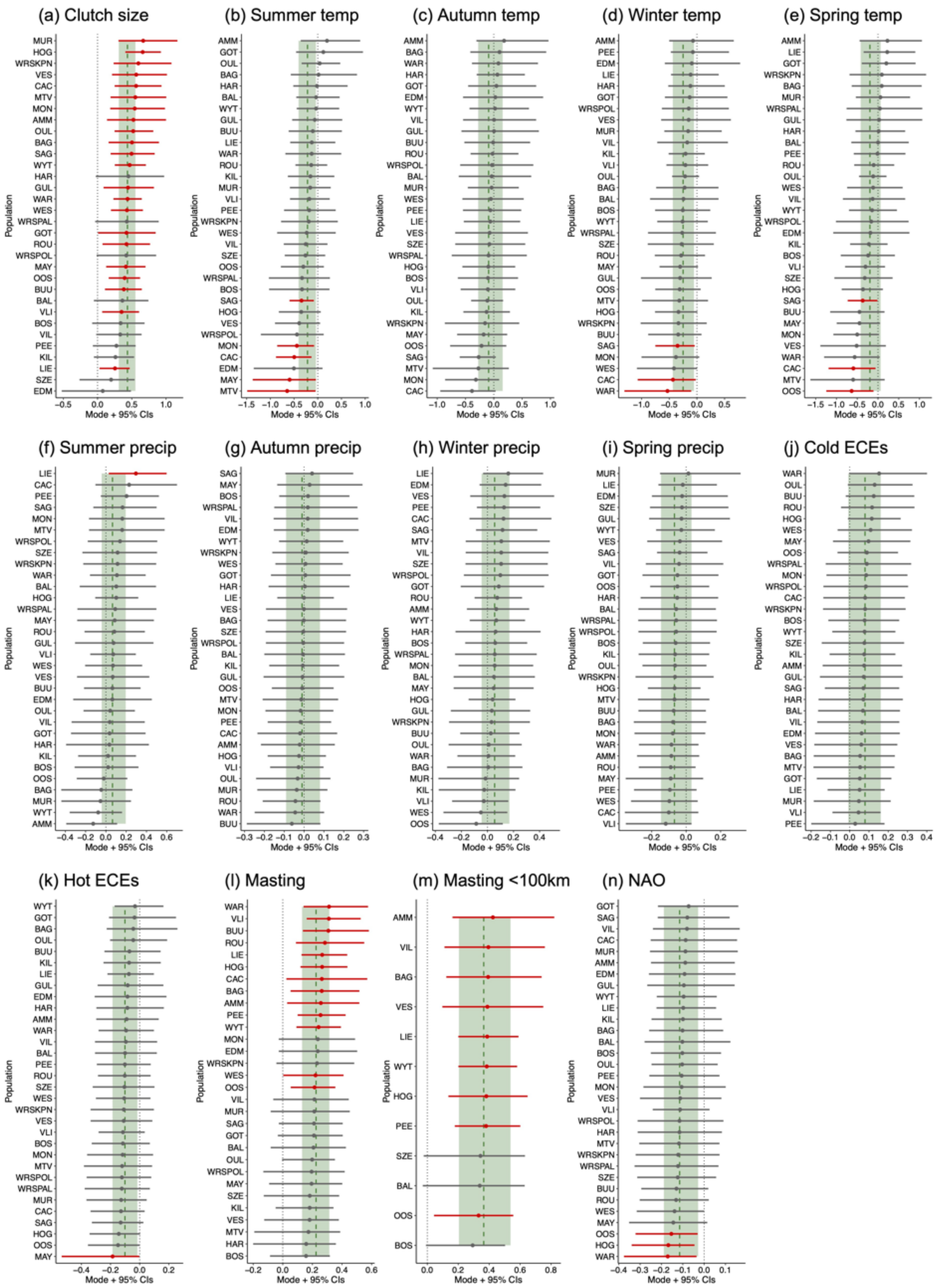


Figure S7 – Population-specific posterior means obtained from linear mixed-effects models which analyse the association between temporal variation in the proportion of breeding juveniles and 14 reproductive and environmental variables (a–n). Each point corresponds to a specific population (on the y-axis), and error bars denote 95% credible intervals. Red points and error bars denote significant effects where credible intervals do not overlap zero (black dashed line). Dotted green line and shading are the overall posterior mode and 95% credible intervals across all populations (as shown in the main text Figure 2).

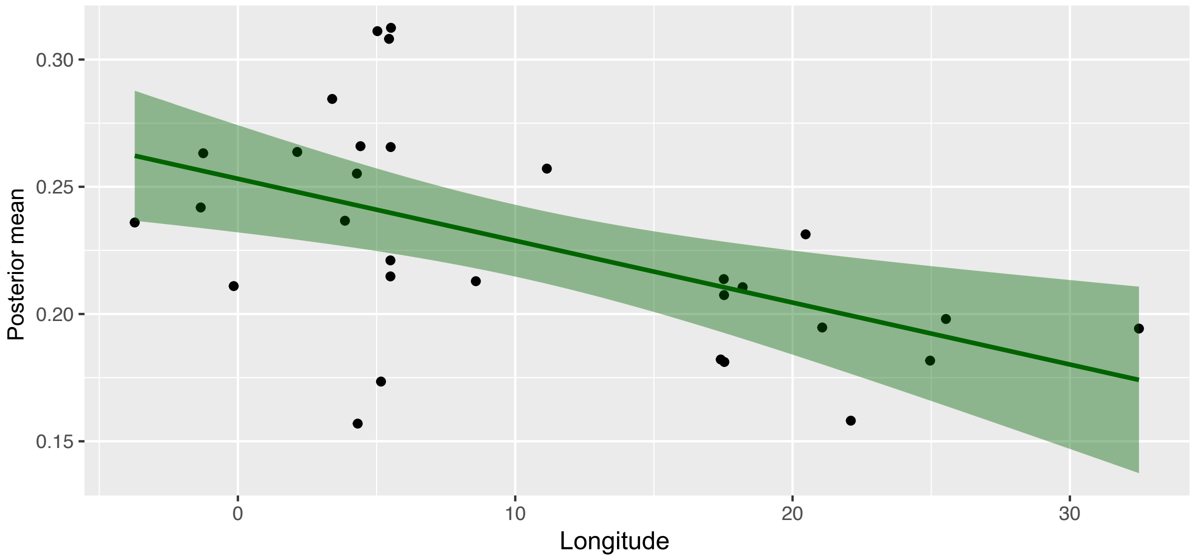


Figure S8 – Relationship between beech mast value and the proportion of breeding juvenile great tits (posterior means from linear mixed-effect model) along the longitude of a population. The line is a linear regression which models the relationship between these two variables, and the shading around this shows the 95% confidence intervals. The relationship is significant and negative (Spearman's correlation: $r$ = -0.517, p = 0.003).

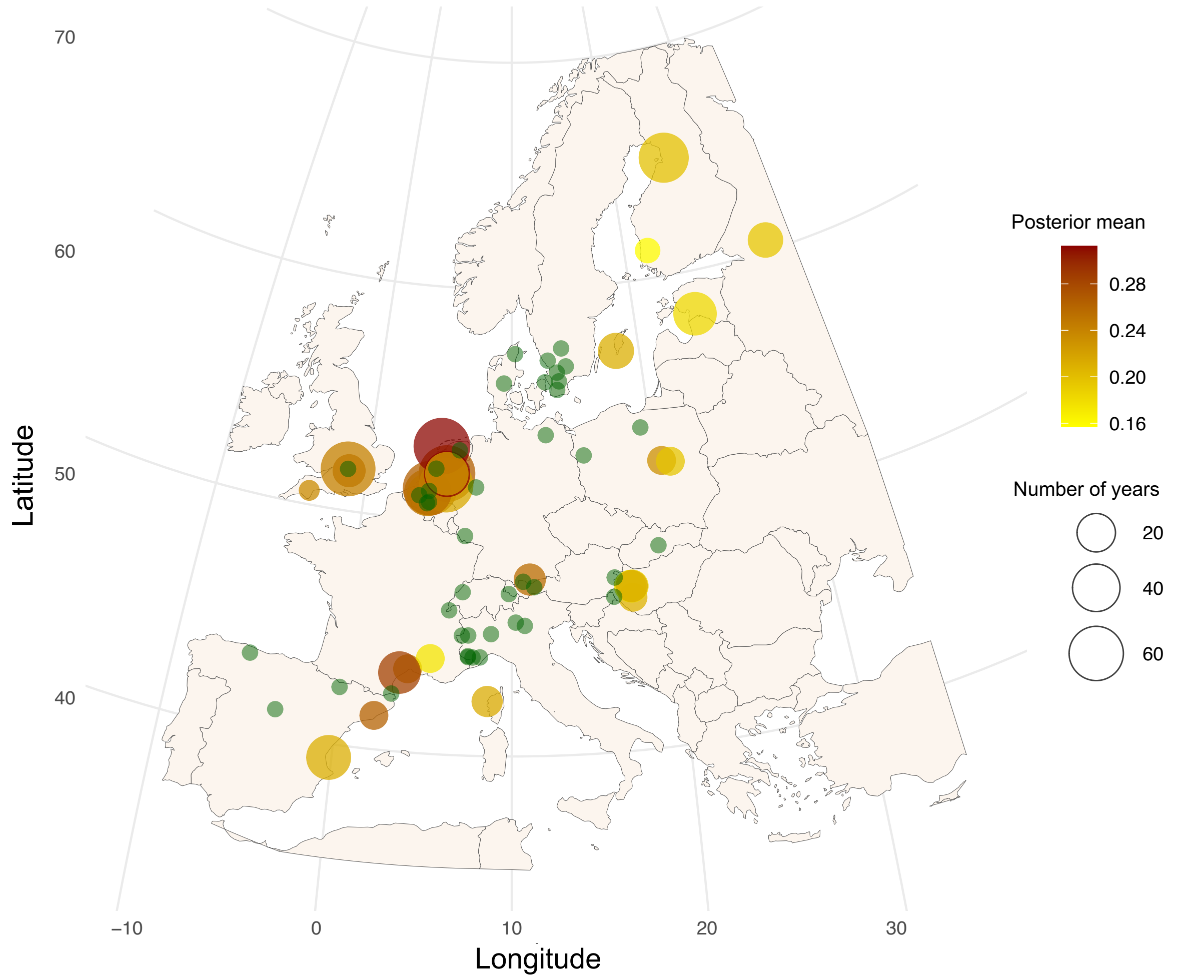


Figure S9 – Map of the 32 great tit study populations across Europe, with point size relative to the number of years in the time series and colour on a scale of yellow to dark red to indicate the strength of association between the proportion of breeding juveniles and beech mast value (the magnitude of the posterior mean obtained from a linear mixed-effects model). The small green points refer to the locations at which mast data was collected and used in this analysis.

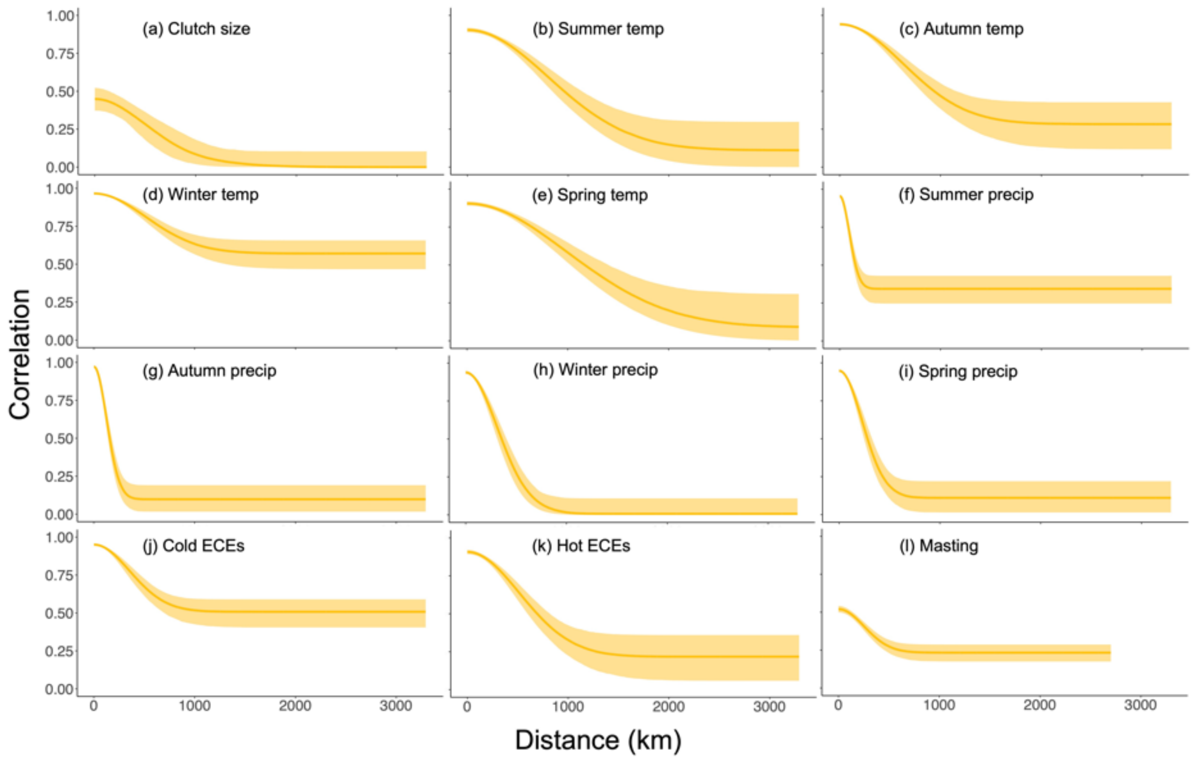


Figure S10 – Spatial synchrony of the reproductive and environmental variables in relation to distance between sites of data collection (i.e. the site of the 32 great tit populations for all variables except beech mast data). In all plots, distance between sites (km) is on the x-axis and correlation between paired sites is on the y-axis. The yellow line is the median estimate of spatial synchrony (calculated in Equation 2 in the main text) based on 2000 bootstrap replicates, with light yellow shading representing 95% credible intervals.

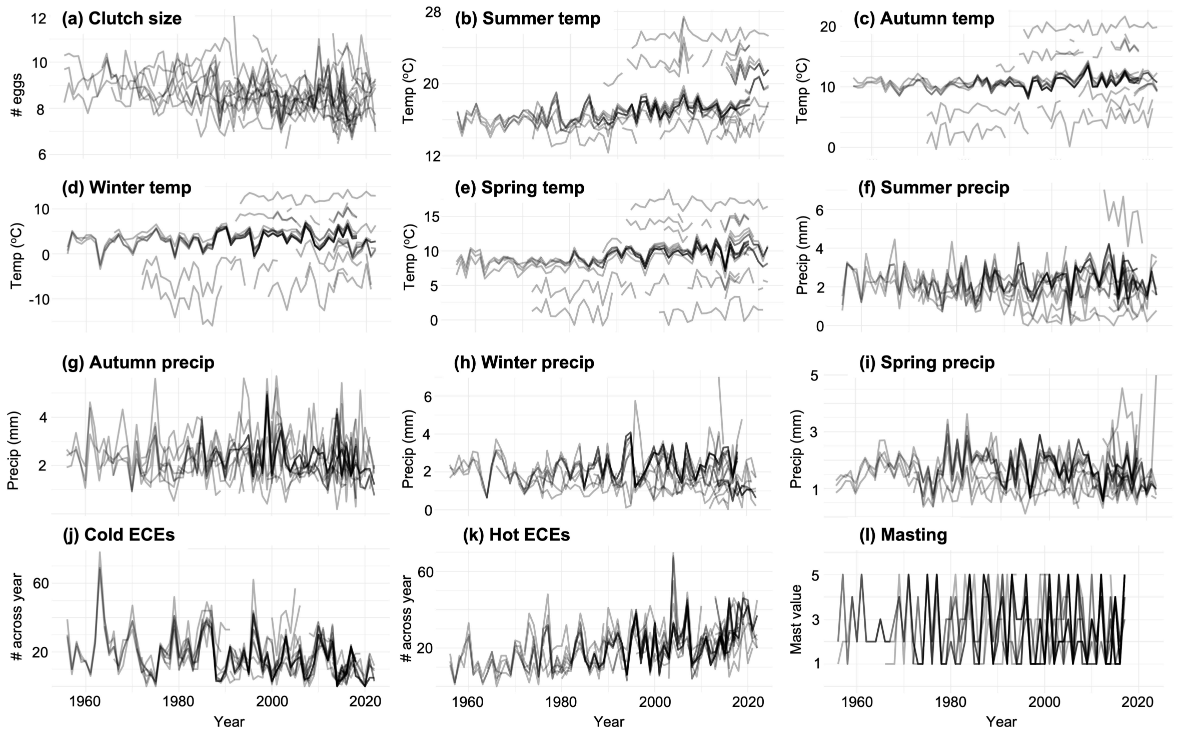


Figure S11 – Local temporal variation in the assessed reproductive and environmental variables for each population of great tits assessed in this study. Figure S3 – Temporal variation in breeding population age structure across European great tit populations. Each line corresponds to a continuous time series for a single population.

### References


Ahola, M.P., Laaksonen, T., Eeva, T. & Lehikoinen, E. (2009). Great tits lay increasingly smaller clutches than selected for: A study of climate- and density-related changes in reproductive traits. *Journal of Animal Ecology*, 78, 1298–1306.

Bailey, L.D. & van de Pol, M. (2016). Tackling extremes: Challenges for ecological and evolutionary research on extreme climatic events. *Journal of Animal Ecology*, 85, 85–96.

van Balen, J.H. (1980). Population fluctuations of the Great Tit and feeding conditions in winter. *Ardea*, 55, 143–164.

Bejer, B. & Rudemo, M. (1985). Fluctuations of Tits (Paridae) in Denmark and Their Relations to Winter Food and Climate. *Ornis Scandinavica*, 16, 29–37.

Bogdziewicz, M., Hacket-Pain, A., Ascoli, D. & Szymkowiak, J. (2021). Environmental variation drives continental-scale synchrony of European beech reproduction. *Ecology*, 102, 1–10.

Bolte, A., Czajkowski, T. & Kompa, T. (2007). The north-eastern distribution range of European beech - a review. *Forestry*, 80, 413–429.

Bordjan, D. & Tome, D. (2014). Rain may have more influence than temperature on nest abandonment in the great tit parus major. *Ardea*, 102, 79–85.

Bouwhuis, S., Sheldon, B.C., Verhulst, S. & Charmantier, A. (2009). Great tits growing old: Selective disappearance and the partitioning of senescence to stages within the breeding cycle. *Proceedings of the Royal Society B: Biological Sciences*, 276, 2769–2777.

Boyce, M.S. & Perrins, C.M. (1987). *Optimizing Great Tit Clutch Size in a Fluctuating Environment*.

Clobert, J., Perrins, C.M., McCleery, R.H. & Gosler, A.G. (1988). Survival Rate in the Great Tit Parus major in Relation to Sex, Age, and Immigration Status. *Journal of Animal Ecology*, 57, 287–306.

Cornes, R.C., van der Schrier, G., van den Besselaar, E.J.M. & Jones, P.D. (2018). An Ensemble Version of the E-OBS Temperature and Precipitation Data Sets. *Journal of Geophysical Research: Atmospheres*, 123, 9391–9409.

Culina, A., Adriaensen, F., Bailey, L.D., Burgess, M.D., Charmantier, A., Cole, E.F., *et al.* (2021). Connecting the data landscape of long-term ecological studies: The SPI-Birds data hub. *Journal of Animal Ecology*, 90, 2147–2160.

Gordo, O. & Sanz, J.J. (2010). Impact of climate change on plant phenology in Mediterranean ecosystems. *Glob Chang Biol*, 16, 1082–1106.

Greño, J.L., Belda, E.J. & Barba, E. (2007). Influence of temperatures during the nestling period on post-fledging survival of great tit Parus major in a Mediterranean habitat. *J Avian Biol*, 0, 071202183307007–0.

Hacket-Pain, A., Foest, J.J., Pearse, I.S., LaMontagne, J.M., Koenig, W.D., Vacchiano, G., *et al.* (2022). MASTREE+: Time-series of plant reproductive effort from six continents. *Glob Chang Biol*, 28, 3066–3082.

Hurrell, J. & Phillips, A. (2003). *NAO Index Data provided by the Climate Analysis Section, NCAR, Boulder, USA, Hurrell*. Available at: https://climatedataguide.ucar.edu/climate-data/hurrell-north-atlantic-oscillation-nao-index-station-based. Last accessed 25 March 2024.

Hurrell, J.W. (1995). Decadal trends in the North Atlantic oscillation: Regional temperatures and precipitation. *Science (1979)*, 269, 676–679.

Julliard, R., McCleery, R.H., Clobert, J. & Perrins, C.M. (1997). Phenotypic adjustment of clutch size due to nest predation in the Great Tit. *Ecology*, 78, 394–404.

Kelly, D. (1994). The evolutionary ecology of mast seeding. *Trends Ecol Evol*, 9, 465–470.

Lamb, P.J. & Peppler, R.A. (1987). North Atlantic oscillation: concept and an application. *Bull. Am. Meteorol. Soc.*, 68, 1218–1225.

Marrot, P., Garant, D. & Charmantier, A. (2017). Multiple extreme climatic events strengthen selection for earlier breeding in a wild passerine. *Philosophical Transactions of the Royal Society B: Biological Sciences*, 372.

Møller, A.P., Balbontín, J., Dhondt, A.A., Adriaensen, F., Artemyev, A., Bańbura, J., *et al.* (2020). Interaction of climate change with effects of conspecific and heterospecific density on reproduction. *Oikos*, 129, 1807–1819.

Moreno, J. & Møller, A.P. (2011). Extreme climatic events in relation to global change and their impact on life histories. *Curr Zool*, 57, 375–389.

Perrins, C.M. (1965). Population fluctuations and clutch-size in the great tit, Parus major L. *Journal of Animal Ecology*, 34, 601–647.
Perrins, C.M. & Moss, D. (1975). Reproductive Rates in the Great Tit. *Journal of Animal Ecology*, 44, 695–706.
Pettifor, R.A., Perrins, C.M. & McCleery, R.H. (2001). The individual optimization of fitness: variation in reproductive output, including clutch size, mean nestling mass and offspring recruitment, in manipulated broods of great tits Parus major . *Journal of Animal Ecology*, 70, 62–79.
Post, E. & Stenseth, N.C. (1999). Climatic variability, plant phenology, and northern ungulates. *Ecology*, 80, 1322–1339.
Regan, C.E. & Sheldon, B.C. (2023). Phenotypic plasticity increases exposure to extreme climatic events that reduce individual fitness. *Glob Chang Biol*, 29, 2968–2980.
Sæther, B.-E., Engen, S., Grøtan, V., Fiedler, W., Matthysen, E., Visser, M.E., *et al.* (2007). The extended Moran effect and large-scale synchronous fluctuations in the size of great tit and blue tit populations. *Journal of Animal Ecology*, 76, 315–325.
Schneider, D.P., Deser, C., Fasullo, J. & Trenberth, K.E. (2013). Climate data guide spurs discovery and understanding. *Eos, Transactions American Geophysical Union*, 94, 121–122.
Silvertown, J.W. (1980). The evolutionary ecology of mast seeding in trees. *Biological Journal of the Linnean Society*, 14, 235–250.
Vacchiano, G., Hacket-Pain, A., Turco, M., Motta, R., Maringer, J., Conedera, M., *et al.* (2017). Spatial patterns and broad-scale weather cues of beech mast seeding in Europe. *New Phytologist*, 215, 595–608.
Wanner, H., Brönnimann, S., Casty, C., Gyalistras, D., Luterbacher, J., Schmutz, C., *et al.* (2001). North Atlantic Oscillation - concepts and studies. *Surv Geophys*, 22, 321–382.
